# Multi-omics analyses of early-onset familial Alzheimer’s disease and Sanfilippo syndrome zebrafish models reveal commonalities in disease mechanisms

**DOI:** 10.1101/2023.10.31.564867

**Authors:** Karissa Barthelson, Rachael A Protzman, Marten F Snel, Kim Hemsley, Michael Lardelli

**Affiliations:** Childhood Dementia Research Group. College of Medicine & Public Health. Flinders Health and Medical Research Institute. Flinders University. Sturt Road, Bedford Park SA 5042 Australia; Alzheimer’s Disease Genetics Laboratory. School of Molecular and Biomedical Sciences, Faculty of Sciences, Engineering and Technology, The University of Adelaide. North Terrace Campus. Adelaide SA 5005 Australia; Proteomics, Metabolomics and MS-Imaging Facility. South Australian Health and Medical Research Institute. North Terrace, Adelaide SA 5000 Australia; School of Physics, Chemistry and Earth Science, Faculty of Sciences, Engineering and Technology, The University of Adelaide. North Terrace Campus. Adelaide SA 5005 Australia

**Keywords:** Alzheimer’s disease, Sanfilippo syndrome, childhood dementia, RNA-seq, zebrafish, extracellular matrix, lysosome, ribosome, oxidative phosphorylation

## Abstract

Sanfilippo syndrome (mucopolysaccharidosis type III, MPSIII) causes childhood dementia, while Alzheimer’s disease is the most common type of adult-onset dementia. There is no cure for either of these diseases, and therapeutic options are extremely limited. Increasing evidence suggests commonalities in the pathogenesis of these diseases. However, a direct molecular-level comparison of these diseases has never been performed. Here, we exploited the power of zebrafish reproduction (large families of siblings from single mating events raised together in consistent environments) to conduct sensitive, internally controlled, comparative transcriptome and proteome analyses of zebrafish models of early-onset familial Alzheimer’s disease (EOfAD, *psen1*^Q96_K97del/+^) and MPSIIIB (*naglu*^A603fs/A603fs^) within single families. We examined larval zebrafish (7 days post fertilisation), representing early disease stages. We also examined the brains of 6-month-old zebrafish, which are approximately equivalent to young adults in humans. We identified substantially more differentially expressed genes and pathways in MPS III zebrafish than in EOfAD-like zebrafish. This is consistent with MPS III being a rapidly progressing and earlier onset form of dementia. Similar changes in expression were detected between the two disease models in gene sets representing extracellular matrix receptor interactions in larvae, and the ribosome and lysosome pathways in 6-month-old adult brains. Cell type-specific changes were detected in MPSIIIB brains at 6 months of age, likely reflecting significant disturbances of oligodendrocyte, neural stem cell, and inflammatory cell functions and/or numbers. Our ‘omics analyses have illuminated similar disease pathways between EOfAD and MPS III indicating where efforts to find mutually effective therapeutic strategies can be targeted.

## Background

Alzheimer’s disease (AD) is the most common form of dementia [1], and is an increasing economic and social burden as societies age. Since the earliest descriptions of AD in the early 1900’s [2], we still do not have effective pharmaceutical treatments which can halt cognitive decline in the long term. This reflects continuing uncertainty regarding the fundamental cause(s) of AD pathology [3, 4].

Most AD cases show late-onset (LOAD, after 65 years of age), and occur sporadically with no evidence of familial inheritance. Less commonly, AD can occur in younger individuals and may be linked to dominant mutations i.e. early-onset familial AD (EOfAD). These dominant mutations occur in the genes presenilin 1 (*PSEN1*) [5], presenilin 2 (*PSEN2*) [6, 7], amyloid β precursor protein (*APP*) [8], or sortilin-related receptor 1 (*SORL1*) [9, 10].

It is clear that the endo-lysosomal system plays a role in the pathogenesis of AD. All the EOfAD genes express proteins with involvement in the cell’s endo-lysosomal system [11–14]. Additionally, many of the genes associated with increased risk for the development of LOAD show endo-lysosomal system involvement [15]. In humans, enlarged endosomes are one of the earliest pathologies observed in AD progression [16]. However, it is difficult to dissect the mechanisms which underlie endo-lysosomal dysfunction in AD, particularly *in vivo*. Damage to the brain observed in post-mortem AD brain tissue is extensive [17]. Many of the lesions seen may be secondary responses to pathological processes so that the information gained may not be directly relevant to therapeutic interventions. Pathological processes which occur early during disease development are likely the most responsive to therapeutic intervention. However, we cannot easily access early diseased human brain tissue for detailed molecular analyses.

Experiments performed in human cell lines can give some insight into the molecular mechanisms underlying AD pathogenesis. However, the relevance of any such observations must be tested *in vivo* (i.e., in animal models), since cultured cells exist in highly artificial environments and some AD-relevant phenomena may be emergent properties of complex multicellular systems such as the brain. A large number of different AD animal models have been developed to investigate AD pathological mechanisms *in vivo* (reviewed in [18, 19]), although the quality/relevance of these models is debated [20, 21]. Much of this uncertainty arises from questions regarding the pathological mode of action of the mutant genes themselves (discussed in [22]). An alternative approach to dissecting the role of the endo-lysosomal system in neurodegeneration is to examine extreme forms of endo-lysosomal system dysfunction, as represented by lysosomal storage disorders that result in childhood dementia.

Sanfilippo syndrome is a type of childhood dementia and has an incidence of approximately 0.17–2.35 per 100,000 live births [23]. It results from one of four recessively inherited lysosomal storage disorders (mucopolysaccharidosis type III, or MPS IIIA, B, C, D; OMIM: 252900, 252920, 252930, 252940, respectively). Like AD, there are no effective therapies are currently approved to treat children with MPS III. However, the genetics of MPS III and the primary pathological mechanisms underlying the disorder are far better understood. MPS III is caused by recessive mutations in genes encoding lysosomal enzymes required for catabolism of the mucopolysaccharide heparan sulfate: N-sulfoglucosamine sulfohydrolase (*SGSH*) [24], N-acetyl-alpha-glucosaminidase (*NAGLU*) [25], heparan-alpha-glucosaminide N-acetyltransferase (*HGSNAT*) [26] and N-acetylglucosamine-6-sulfatase (*GNS*) [27]. Incomplete catabolism of heparan sulfate (HS) leads to its accumulation in lysosomes. Mutations in each of these genes define subtypes of Sanfilippo syndrome, i.e. mutations in *SGSH*, *NAGLU*, *HGSNAT*, and *GNS* define MPS IIIA, IIIB, IIIC, and IIID respectively. Arylsulfatase G (*ARSG*) deficiency causing MPS IIIE has been observed in mice [28]. *ARSG* mutations in humans are associated with Usher syndrome – a condition associated with hearing and vision loss. However, it is debated whether this disease should be classified as MPS IIIE (reviewed in [29]).

While AD and MPS III are distinct disorders, MPS III is often colloquially referred to as “childhood Alzheimer’s”, due to the similarity to AD of symptoms experienced by children living with MPS III. At the cellular level in the brain, there are commonalities of phenomena such as oxidative stress [30, 31], inflammation [31–33], amyloid β and tau-related pathological processes [34–36], and, importantly, dysfunction of the endo-lysosomal system [37]. However, these two diseases have never been compared directly within one experimental system.

Zebrafish genetic models have emerged as powerful tools for studying cellular/molecular phenomena in neurodegenerative diseases, particularly due to their ∼70% genetic similarity to humans [38, 39], similarity in brain architecture (reviewed in [40]) and their high fecundity (reviewed in [41]). We have previously generated several zebrafish knock-in models of mutations causing early-onset familial AD [42–46] and MPS III [47] and have analysed their brain transcriptomes. We have noted that all the models of both diseases exhibit differential expression of genes involved in metabolic pathways [47, 48].

The most sensitive comparisons of these disease models are best done within single families. This is because family-specific effects reduce statistical power to detect changes in the phenotype of interest [49]. Within single families, genetic and environmental variation is reduced between experimental groups of interest, allowing the detection of subtle changes to gene expression. This approach is widely utilised in transcriptomics in zebrafish [50–53]. Here, we use intra-family analyses to compare the transcriptomes and proteomes of a zebrafish model of EOfAD; *psen1*^Q96_K97del^ [54], with those of *naglu*^A603fs^, a model of MPS IIIB [47]. We performed our analyses at two ages: 7 days post fertilisation (dpf) which models very early, pre-symptomatic stages for both diseases, and in young adult brains (6 months of age), which models pre-symptomatic disease stages for EOfAD but is a symptomatic stage in MPS IIIB zebrafish [47]. We identified distinct changes in gene expression and protein abundances in each model at both ages. However, important age-related commonalities are also observed in genes implicating the extracellular matrix, mitochondria, lysosomes and ribosomes, indicating fundamental, underlying similarities in pathological mechanisms.

## Methods

### Zebrafish husbandry and breeding strategy

The generation of the *naglu*^A603fs^ and *psen1*^Q96_K97del^ lines of zebrafish used in this study was described in [47] and [54] respectively. All fish were maintained and raised in a recirculating water system with a constant temperature of approximately 26°C, salinity levels at approximately 300 ppm, and conductivity approximately 800 µS/cm. These parameters were measured using an ExStik EC500 meter (Extek Instruments, Nashua, New Hampshire, United States). The pH level was measured using pH strips (Sigma Aldrich, St. Louis, Missouri, United States) and was maintained between 6 and 7. All fish were raised under a 14:10 hour light:dark cycle. Fish were fed twice daily with a diet of NRD 5/8 dry food (Inve Aquaculture, Dendermonde, Belgium) in the mornings and live *Artemia salina* (Inve Aquaculture) in the afternoon.

### Harvesting of samples for RNA-seq and proteomics

To generate the zebrafish samples for the larval RNA-seq experiment, we crossed a *psen1*^Q96_K97del/+^; *naglu*^A603fs/+^ zebrafish with a *psen1^+/+^*; *naglu*^A603fs/+^ zebrafish, to generate a family of 100 sibling larvae with a variety of genotypes (**Fig.1**). These larvae were raised together in a 14 cm diameter plastic petri dish (VWR, Radnor, Pennsylvania, United States) in 150 mL of E3 medium [55] (pH ∼7) until 7 days post fertilisation (dpf). The E3 medium was changed every two days.

**Fig. 1:**
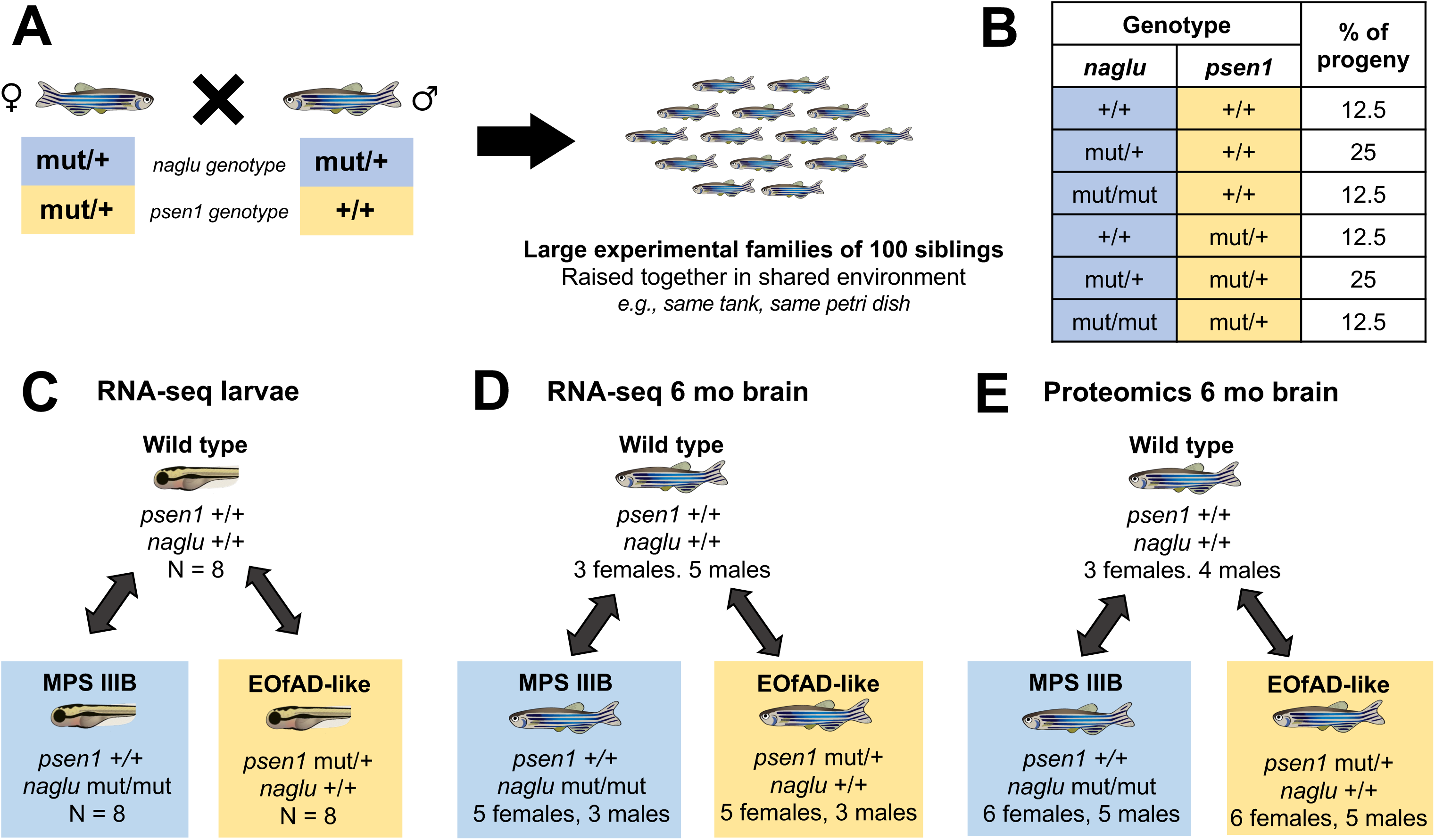
Intra-family transcriptome analysis strategy. **A)** A female zebrafish heterozygous for both the *naglu*^A603fs^ (MPS IIIB) and the *psen1*^Q96_K97del^ (EOfAD-like) mutations was mated with a male zebrafish only heterozygous for *naglu*^A603fs^. The subsequent family of ∼100 sibling zebrafish are raised together in a shared environment. **B)** Table showing expected genotype proportions in the MPS IIIB/EOfAD-like families. **C)** Schematic showing the pairwise comparisons between MPS IIIB and EOfAD-like zebrafish to wild type zebrafish in each RNA-seq experiment for each age. Detailed numbers of number of fish per genotype and sex can be found in **Fig.S1**.

At 7 dpf, 100 sibling larvae were euthanised by cold shock in ice water, then individual larvae were placed in 100 µL of RNA*later*™ Stabilization Solution (Invitrogen™, Waltham, Massachusetts, United States). The larvae in RNA*later* were incubated overnight at 4°C. Then, the tails of the RNA*later-*preserved larvae were removed for genotyping using sterile needles to cut transversely at the level of the cloaca. The anterior ends of the larvae were stored at −80°C until use (**Fig.S1**).

For analysis of 6-month-old adult brains, we raised two additional families of zebrafish spawned by different parental pairs (**Fig.S1**). All fish in the families (at least 100 fish per family) were euthanised by ice cold water shock, the sex of each fish was confirmed by dissection and observation of testes or ovaries, entire heads were separated from bodies at the level of the gills, and tails were removed for genotyping by PCR.

For RNA-seq analysis, each entire head was immediately placed in 600 µL of RNA*later* solution and incubated at 4°C overnight. Subsequently, the RNA*later*-preserved brain was dissected from the skull, and then stored in a fresh aliquot of 100 µL of RNA*later* at −80°C until analysis. For proteomics analysis, the brain was dissected from the skull within two minutes of death, snap frozen in liquid nitrogen, and stored at −80°C until use.

### PCR genotyping

Genomic DNA (gDNA) was isolated from the tails of 7 dpf larvae and 6-month-old adults by boiling in sodium hydroxide as described in [55]. These gDNA preparations were then used as templates for genotyping by polymerase chain reactions (PCRs). Our method of genotyping involved PCR amplification across mutation sites using the primers described in **Table 1** (purchased from Sigma Aldrich). Then, we subjected the amplified PCR products to electrophoresis using gels consisting of 2-3% agarose (Sigma Aldrich) in 1 x Tris-acetate-EDTA (1 x TAE) buffer for 2 hours at 90V, alongside DNA marker ladders generated from the pHAPE plasmid [56]. In this way, we could resolve two PCR products, one each resulting from the wild type allele and/or mutant allele of each gene (see **Fig.S2,S3**).

**Table 1:**
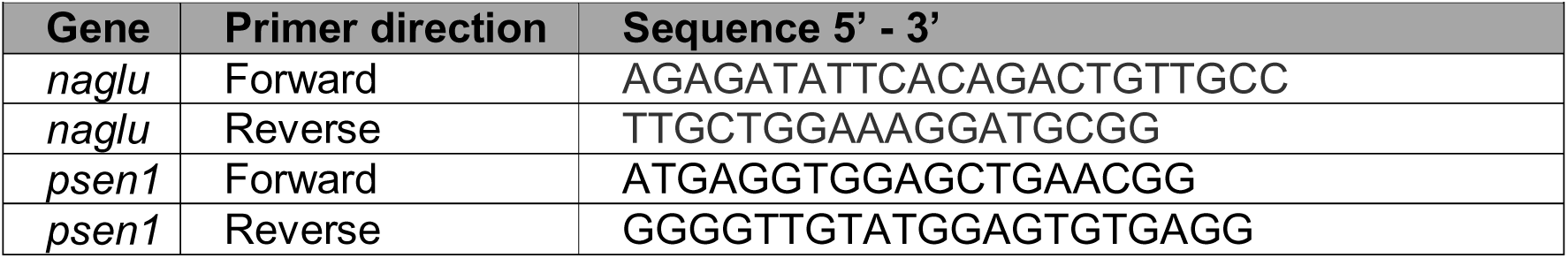
Genotyping primer details.

### RNA extraction and sequencing

Total RNA was isolated from the individual larval zebrafish and 6-month-old adult brains using the *mir*Vana™ miRNA Isolation Kit (Invitrogen) following the manufacturer’s instructions. Only fish with genotypes *psen1* ^+/+^; *naglu* ^+/+^ (wild type), *psen1* ^Q96_K97del/+^; *naglu ^+/+^* (EOfAD-like) and *psen1 ^+/+^*; *naglu ^A603fs/A603fs^* (MPS IIIB) were analysed as these represent the genetic states of the respective human diseases. In the larval RNA-seq experiment, we analysed n = 8 larvae per genotype (**Fig.1C**). For analysis of adult brain transcriptomes, we analysed at least n = 3 per genotype and sex: 3 female and 5 male wild type brains, 5 female and 3 male MPS IIIB brains, and 5 female and 3 male EOfAD-like brains (**Fig.1D**). To remove any genomic DNA, we treated the total RNA isolates with DNase I using the *DNA-free™* DNA Removal Kit (Invitrogen) following the manufacturer’s instructions for routine DNase digestion. Then, 400 ng of total RNA from larva or adult brain were then delivered to the South Australian Genomics Center (SAGC, Adelaide, South Australia, Australia) for stranded polyA+ library preparation using Nugen Universal Plus mRNA-seq (Tecan, Männedorf, Switzerland), protocol M01442 v2. The libraries were further processed using the MGIEasy Universal Library Conversion Kit, Part No. MGI1000004155 (MGI, Shenzhen, China), then sequenced (2 x 98 paired-end with 8 bp unique molecular identifiers) using the MGI DNBSEQ-G400 platform across four lanes which were subsequently merged.

### RNA-seq data pre-processing

We pre-processed the raw fastq files using a custom pipeline implemented in *snakemake* [57]. For details of this pipeline, see **Fig.S4** and https://github.com/karissa-b/2021_MPSIIIBvQ96-RNAseq-7dpfLarve. Briefly, *fastp* [58] was used to trim adaptors and filter the reads by quality and length. Then, the reads were aligned to the zebrafish genome (GRCz11, Ensembl release 104) using *STAR* [59] to generate sorted .bam files. These were subsequently indexed using *samtools* [60]. After alignment, PCR duplicates were de-duplicated using the *dedup* function of *umi-tools* [61]. The number of reads aligning to gene models of the GRCz11 genome were counted using *featureCounts* [62]. *FastQC* [63] and *ngsReports* [64] were used to assess the quality of the raw and trimmed data, as well as the original and deduplicated alignments.

### Statistical analysis of RNA-seq data

Statistical analysis of the RNA-seq data was performed using R [65]. We first omitted lowly expressed genes, which are statistically uninformative for differential gene expression analysis. We defined a gene as lowly expressed if it contained a log counts per million (logCPM) of less than 0.5. To account for genes expressed below this limit of detection in some experimental groups, but not in the others, we retained the genes which fell below this threshold if they contained sufficient logCPM values in at least 8 out of the 24 total RNA-seq samples (i.e. one of the mutant genotype groups) in each experiment for each age.

Differential gene expression analysis was performed with likelihood ratio tests using *edgeR* [66]. At 7 dpf, zebrafish larvae, are yet to become sexually dimorphic and so we specified a design matrix with the wild type genotype as the intercept, and each of the mutant genotypes as coefficients. For 6-month-old adult brains, we specified an intercept-free design matrix with each genotype and sex combination as the coefficients (e.g. ∼0 + group). This was to allow pairwise contrasts between each mutant and their wild type siblings within sexes. We considered a gene to be differentially expressed (DE) if the p-value after adjusting for the false discovery rate (FDR) was less than 0.05.

We observed biases for differential expression of genes due to their guanine-cytosine content (%GC) and their length (**Fig.S5**). To account for this bias, we used conditional quantile normalisation (*cqn*, [67]). The offset term generated by *cqn* was then included in an additional differential gene expression analysis with *edgeR* to give the final list of DE genes.

Over-representation of gene ontology (GO) terms [68, 69] within the DE genes was identified using *goseq* [70], setting a %GC content measure as a co-variate for each gene. This measure was calculated as the average %GC content per transcript, weighted by the transcript length, to account for the fact that different alternative transcripts per gene will have different %GC contents. We considered a GO term to be significantly over-represented in the DE genes if the FDR-adjusted p-value from *goseq* was < 0.05.

Due to the highly overlapping nature of GO terms, and the limitation of the hard threshold of considering a gene to be DE, we also performed gene set enrichment analysis of the KEGG [71] pathway gene sets using an ensemble approach. We calculated the statistical significance of the changes to gene expression in the KEGG pathways using three gene set enrichment tools, each having different statistical methodologies: *fry* [72], *camera* [73] and *GSEA* [74, 75] as described previously [44]. Then, the individual p-values were combined using the harmonic mean p-value (HMP) method, which has been shown specifically to be robust to dependent tests [76]. To further protect from type I errors, we considered a gene set to be significantly altered if the FDR-adjusted HMP was < 0.05.

To assess the statistical power achieved in these experiments, we used the *ssize_vary*() function of *ssizeRNA* [77] using default parameters. Briefly, we defined the number of genes considered detectable, the average read counts (accounting for the offset by *cqn*) in the wild type group, and the dispersion estimate from each experiment. The proportion of non-DE genes was set to 0.9, the pseudo-sample size to 30, the FDR level to 0.05, and the range of absolute logFC values from 0-2.

### Proteomics

For brain proteomics analysis, we analysed at least n = 3 fish per genotype and sex: 3 female and 4 male wild type brains, 6 female and 5 male MPS IIIB brains, and 6 female and 5 male EOfAD-like brains. An EasyPrep Micro MS Sample Prep Kit (Part No. A57864, Thermo Scientific) was used for proteomic sample preparation. First, 100 µL of lysis buffer from the kit was added to each Precellys homogenisation tube (Part No. P000933-LYSK0-A.0, Bertin Technologies). Frozen tissue samples were transferred to each tube and homogenised at 4 °C in a Precellys 24 Tissue Homogeniser (Bertin Technologies) at 4000 rpm for 2 rounds of 30 seconds with a 30 second pause between rounds. Pierce BCA Protein Assay (Part No. 23227, Thermo Scientific) was performed to quantify total proteins, and the proteins were then normalised to 10 µg protein and brought up to a total volume of 10 µL in lysis buffer. The sample preparation kit protocol was followed for the digestion and sample cleanup, with the sample being digested for 2.5 hours at 37 °C. After drying down the peptides in a vacuum concentrator, each sample was reconstituted to 200 ng peptides (negating sample loss) in loading buffer (2% MS-grade acetonitrile, 0.1% trifluoroacetic acid).

Prior to mass spectrometric analysis, samples were separated by reversed-phase liquid chromatography (LC) using a Waters ACQUITY UPLC M-Class System equipped with an Acclaim PepMap 100 C18 trap cartridge (5 mm x 1 mm, 5 µm, Part No. 160434, Thermo Scientific). During the 5-minute loading of the sample to the trap cartridge, Pump A of the Auxiliary Sample Manager (ASM) flowed at a rate of 10 µL/min with loading buffer, while the µBinary Solvent Manager flowed at a rate of 0.4 µL/min with 97% mobile phase A (MPA; 0.1% formic acid in water) and 3% mobile phase B (MPB; 0.1% formic acid in acetonitrile). This was followed by chromatographic separation at a constant 0.4 µL/min flow rate using an Aurora ULTIMATE column (25cm x 75µm, 1.7µm C18) fitted with a captive spray ionisation emitter (Part No. AUR3-25075C18-CSI, IonOpticks). The gradient started at 3% MPB, gradually increasing to 17% at 60 min, 25% at 90 min, 37% at 100 min and 95% at 110 min; it was held at 95% MPB until 120 mins to wash the column, sharply dropped to 3% at 121 mins and left to equilibrate until the end of the run at 130 mins.

A timsTOF Pro2 mass spectrometer (Bruker) equipped with a captive spray ionisation source was used to collect data in positive ionisation mode using the dia-PASEF long gradient method with default settings. Raw mass spectrometric data was converted into .htrms files before being uploaded to Spectronaut v. 19.0 (Biognosys) for database searching. Proteins were identified and quantified using Spectronaut DeepDIA without cross-run normalisation and run against the zebrafish proteome FASTA database (Proteome ID: UP000000437), downloaded as one protein sequence per gene, alongside the FASTA file for porcine trypsin (Accession No. P00761) from Uniprot.org.

### Statistical analysis of MS/MS data

Only proteins detected in all samples of genotype and sex were retained. Out of the 9,942 proteins detected in the analysis, 1421 showed missing values and these proteins were omitted from the analysis. The remaining 8521 relative protein abundances were log-transformed and quantile-normalised before being fitted to linear models using *limma* [78]. The design matrix was specified without an intercept, and each genotype and sex combination as the coefficients (e.g., 0 + group). A contrasts matrix was also defined to compare each of the mutant genotypes to their wild type siblings within each sex. We considered a protein to be differentially expressed (DE) in each contrast if the p-value after adjusting for the false discovery rate (FDR) was less than 0.05. Functional analysis of the DE proteins was performed using STRING [79], and using the harmonic mean as described for the transcriptome data.

Visualisation of the transcriptome and proteome data was performed using *ggplot2* [80], *tidyheatmap* [81], *pathview* [82] and *UpSetR* [83].

## Results

### What are the transcriptomic changes shared by MPS IIIB and *psen1* EOfAD-like zebrafish larvae?

We sought to compare directly the changes in global gene expression occurring due to heterozygosity for an EOfAD-like mutation in *psen1* (*psen1*^Q96_K97del^) or homozygosity for loss-of-function mutations in *naglu* (*naglu*^A603fs^, i.e. causing MPS IIIB) in 7 dpf zebrafish larvae using RNA-seq. First, to assess the overall similarity between samples, we used principal component analysis (PCA). No distinct clustering of samples by genotype was observed (**Fig.2A**), supporting that these mutant genotypes do not result in extensive changes to larval zebrafish bulk transcriptomes.

**Fig. 2:**
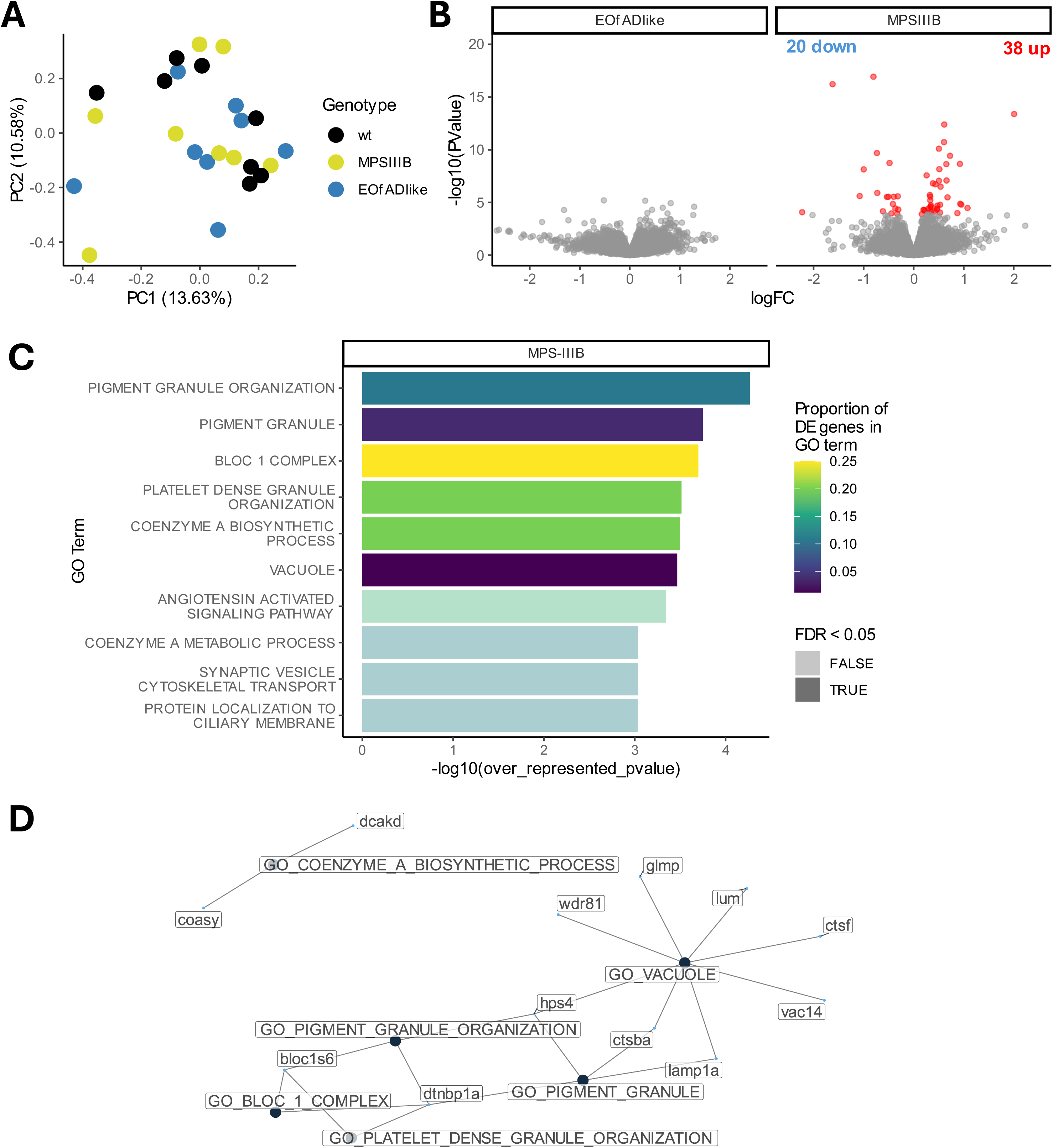
Differential gene expression analysis in zebrafish larvae at 7 days post fertilisation (dpf). **A)** Principal component analysis (PCA) on the log_10_ counts per million (logCPM) values after adjusting for %GC and length bias using cqn [67]. Each point represents a larval zebrafish transcriptome sample which is coloured by genotype. The percentage variations that principal component 1 (PC1) and 2 (PC2) explain in the dataset are indicated in parentheses in the axis labels. **B)** Numbers of differentially expressed genes for each mutant genotype relative to wild type zebrafish larvae. **C)** Volcano plots showing the log fold change (logFC) and negative log_10_ of the p-value of the genes expressed in MPS IIIB and EOfAD-like larval zebrafish samples. Each point represents a gene which is coloured red if the differential expression FDR-adjusted p-value was less than 0.05. The x-axis labels are constrained to within −2 and 2 for visualisation purposes. **D)** The 10 most significantly over-represented Gene Ontology (GO) terms among the differentially expressed genes in MPS IIIB zebrafish larvae. The vertical black line represents where the FDR = 0.05. **E)** Network visualisation indicating the overlap of DE genes within the significantly over-represented GO terms in MPS IIIB larvae. GO term nodes (black) are connected by unweighted edges to the nodes representing DE genes (blue) to which they are assigned.

We next asked whether any genes were differentially expressed (DE) in both MPS IIIB and EOfAD-like zebrafish larvae, which may support some shared pathological changes. However, we did not identify any genes with an FDR-adjusted p-value < 0.05 in the EOfAD-like larvae (meaning that, at the single gene level, it was not possible to identify any similarity in the changes to gene expression in EOfAD-like and MPS IIIB zebrafish larvae). In contrast, 57 DE genes were identified for the MPS IIIB mutant genotype (**Fig.2B** and full output in **Supplemental Table 1**) consistent with the earlier onset and greater disease severity of MPS III relative to EOfAD. The DE genes in the MPS IIIB larvae were significantly over-represented with 8 GO terms (**Fig.2C**, full output in **Supplemental Table 2**) and the DE genes within these significant GO terms were typically linked to multiple GO terms. Consequently, the enrichment of DE genes within these GO terms is influenced to some extent by the same set of genes (**Fig.2D**).

To detect any subtle changes in gene expression, we then took an ensemble approach to gene set testing utilising the calculation of the harmonic mean p-value (in which the input does not rely on pre-defining DE genes). We applied this method to the gene expression data from both mutants, using the KEGG [71] gene sets. In the MPS IIIB larvae we detected significant changes to gene expression in 13 KEGG gene sets (**Fig.3A**, full output in **Supplemental Table 3**). Not surprisingly, the gene sets for *lysosome*, *glycosaminoglycan degradation* and *other glycan degradation* were found to be significantly altered, consistent with the primary disease mechanisms in MPS III (reviewed in [84]) (**Fig.3A** and **Fig.S6)**. Only 1 KEGG gene set was significantly altered in the EOfAD-like mutant larvae; *ECM receptor interaction*. This gene set is also significantly altered in the MPS IIIB larvae. A positive Pearson correlation was found for the logFC of the genes in this gene set between the EOfAD-like and MPS IIIB mutants (**Fig.3B**), and the genes in the pathway tended to be downregulated (**Fig.S7**). These observations suggest that changes to ECM function could be a common, early disease mechanism between EOfAD and MPS III.

**Fig. 3.**
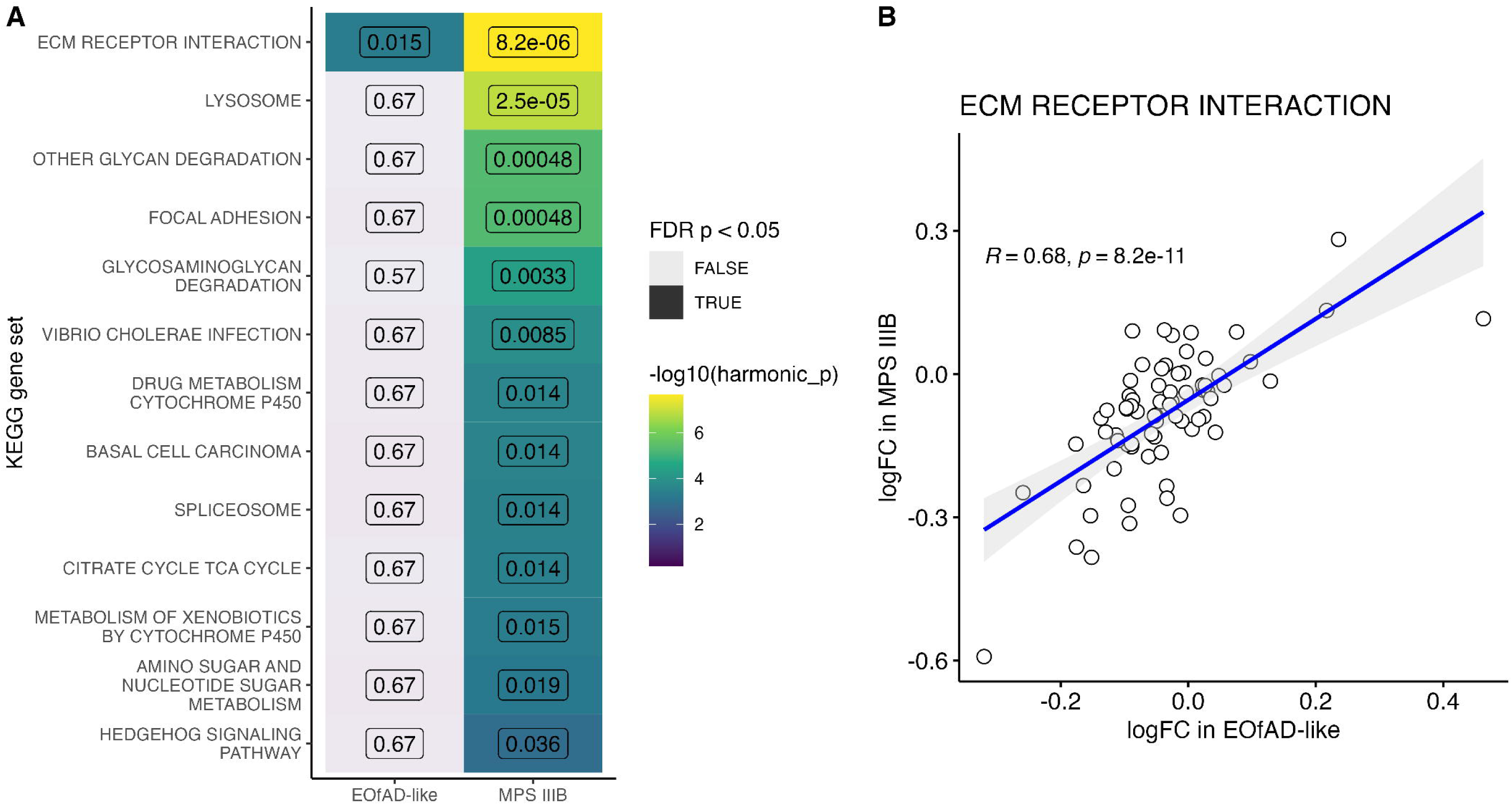
KEGG pathway analysis in zebrafish larvae at 7 days post fertilisation (dpf). **A)** Heatmap indicating the KEGG gene sets significantly altered in at least one of the mutant larval zebrafish samples. Cells in the heatmap are coloured by the log_10_ of the harmonic mean p-value (i.e. the brighter, more yellow cells are more statistically significant). These appear shaded and grey if the p-value did not reach the significance threshold of p < 0.05. The numbers inside the heatmap are the FDR-adjusted harmonic mean p-values. **B)** Pearson correlation of the logFC values of genes in the KEGG gene set for extracellular matrix (ECM) receptor interaction. Grey shading indicates confidence intervals.

We did not detect any differentially expressed genes in the n = 8 individual EOfAD-like mutants (**Fig.2B,C**), while our previous analysis of this EOfAD-like mutation that was performed on n = 6 pools each of 40 larvae at 7 dpf had identified 228 DE genes [85]. Therefore, we hypothesised that our experiment was statistically underpowered for the use of individual larvae. Subsequently, we performed a post-hoc power calculation using *ssizeRNA* [77]. Indeed, based on the variation observed within the wild type genotypes, our experimental structure had achieved an estimated 51% power to detect DE genes. A sample of size of > 20 individual larvae would be required to achieve 70% power (**Fig.S8A**). This is not feasible to achieve within a single zebrafish family as, in our experience, the maximum size for a single clutch of zebrafish rarely exceeds 150 viable embryos. Therefore, we decided to repeat the analysis using adult zebrafish brains, where we have previously shown that n = 6 generally gives approximately 70% power to detect significant changes in gene expression [44].

### What do the young adult brains of MPS IIIB and *psen1* EOfAD-like zebrafish have in common?

We first performed a post-hoc power calculation on the 6 month old adult brain RNA-seq data and confirmed we achieved approximately 80% power to detect DE genes (**Fig.S8B**). We next explored the overall similarity between brain transcriptomes using PCA. This revealed some clustering of samples: the MPS IIIB samples formed a cluster distinct from the wild type and EOfAD-like samples in both sexes (**Fig.4A-C**). This suggests that the 6-month-old MPS IIIB brain transcriptome does not represent a wild type state, and that significant changes are apparent. We next asked which genes were differentially expressed due to MPS IIIB and EOfAD-like genotypes within each sex. Pairwise contrasts between MPS IIIB mutants with their wild type siblings revealed 199 and 352 DE genes in females and males respectively. In contrast, the EOfAD-like mutants exhibited far fewer DE genes: 12 in females and 4 in males (**Fig. 4D**, full output in **Supplemental Data 1**). To determine the biological context of these DE genes, we performed over-representation analysis of gene ontology (GO) terms using *goseq*. Enrichment of GO terms implicating lysosomes (e.g. lysosomal lumen), and oligodendrocytes (e.g. oligodendrocyte differentiation) were identified only in MPS IIIB mutants (full output in **Supplemental Data 2**). No significant over-representation of GO terms were found in the EOfAD-like mutants (**Supplemental Data 2**), likely due to the low number of DE genes in these mutant fish.

**Fig. 4:**
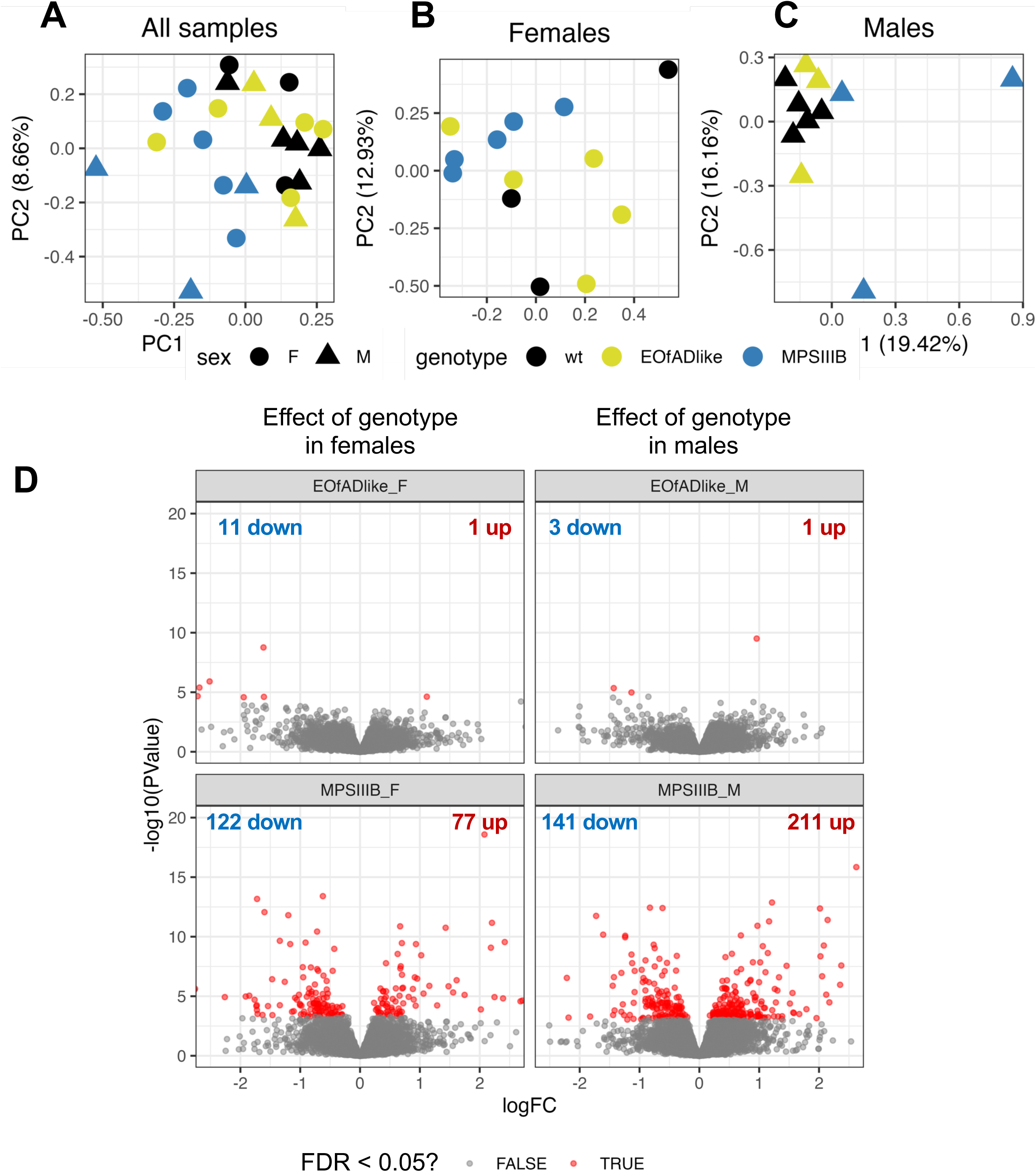
Differential gene expression analysis in zebrafish brains at 6 months of age. Principal component analysis (PCA) on the log_10_ counts per million (logCPM) values after adjusting for %GC and length bias using *cqn* [67]. Each point represents an adult zebrafish brain transcriptome sample in **A)** all samples, **B)** only female samples and **C)** only male samples that are coloured by genotype and shaped by sex (F: female, M: male, wt: wild type). The percent variations that principal component 1 (PC1) and 2 (PC2) explain in the dataset are indicated in parentheses in the axis labels. **D)** Volcano plots showing the log fold change (logFC) and negative log_10_ of the p-value of the genes expressed in MPS IIIB and EOfAD-like zebrafish adult brains. Each point represents a gene which is coloured red if the gene was differentially expressed at a FDR-adjusted p-value of less that 0.05. The x-axis labels are constrained to within −2 and 2 for visualisation purposes.

Due to the low number of DE genes in the EOfAD-like mutants, we next tested the KEGG gene sets using the ensemble approach to gene set enrichment analysis, where pre-defining DE genes is not required. Using this method, we detected 3 gene sets significantly altered in both EOfAD-like and MPS IIIB mutant brains in at least one sex: *lysosome*, *oxidative phosphorylation*, *ribosome* **(Fig.5A**, full output in **Supplemental Table 3)**. All these gene sets were broadly upregulated (**Fig.5B-D**).

**Fig. 5:**
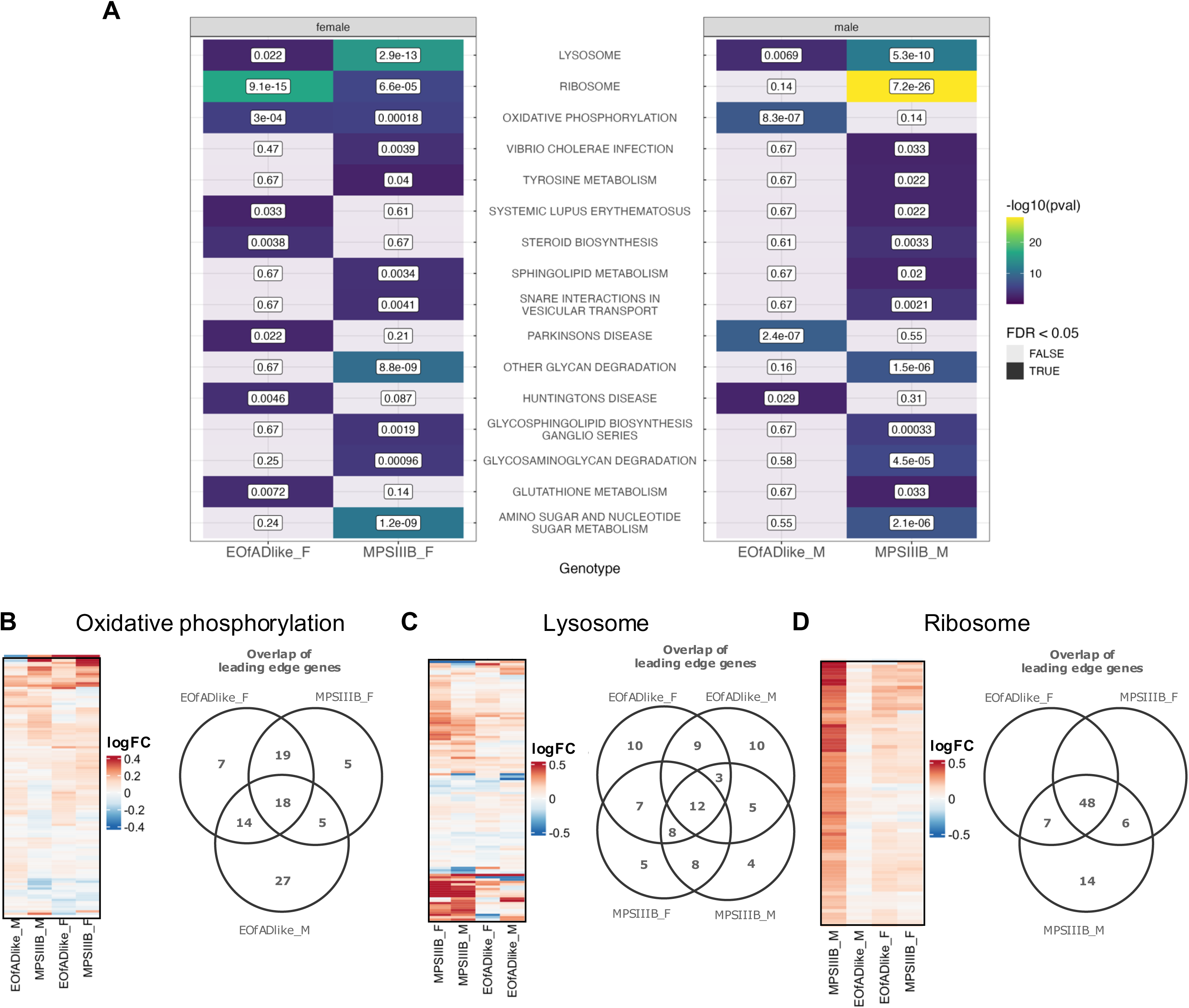
KEGG gene sets significantly altered in zebrafish brain transcriptomes at 6 months of age. **A)** Heatmap indicating those KEGG gene sets significantly altered in least two comparisons of mutant zebrafish to wild type. Cells in the heatmap are coloured by the log_10_ of the harmonic mean p-value (i.e. the brighter, more yellow cells are more statistically significant). They appear grey if the p-value did not reach the significance threshold of < 0.05. The numbers indicate the FDR-adjusted p-value. **B)** Heatmaps indicating the similarities in the log fold change (logFC) of genes in the KEGG gene sets for *lysosome*, **C)** *oxidative phosphorylation* **D)** *ribosome* and **E)** *other glycan degradation* in EOfAD-like and MPS IIIB mutant zebrafish brains. Upset plots display the overlap of leading-edge genes across the gene sets. The numbers next to each contrast represent the total number of leading-edge genes. The numbers above the bars indicate the number of genes shared between the gene sets in each contrast shown in the corresponding intersections.

To assess whether these gene sets were similarly affected in each mutant, we examined the leading edge genes identified by *fgsea* [74], The leading edge genes are those which contribute the most to the enrichment score, and can be interpreted as the core genes driving the statistical significance of the gene set. For the *oxidative phosphorylation* gene set, 37 leading edge genes were shared between female EOfAD-like (**Fig.5B**) and female MPS IIIB brains (**Fig.5B**). This represents more than half of the leading edge genes in each comparison, supporting that this gene set is altered in similar way in these female mutants. Only 18 genes were common across all leading edge genes in MPS IIIB and EOfAD-like mutants, which represents 38% (MPS IIIB females), 31% (EOfAD-like females), and 28% (EOfAD-like males) of the leading edge genes. These observations support that the *oxidative phosphorylation* pathway is partially altered in a similar way in female MPS IIIB and female EOfAD-like mutant zebrafish, and in a different way in EOfAD-like males.

For the *lysosome* gene set, 14 leading-edge genes were shared between both mutants and sexes (**Fig.5C**). In females, a total of 27 leading edge genes were shared between MPS IIIB and EOfAD-like mutants (68% and 54% of the leading edge genes respectively). In males, 20 leading edge genes were shared, representing 51% (EOfAD-like) and 50% (MPS IIIB) of all leading edge genes in each mutant. These results support that the lysosome gene set is only partially similar in its dysregulation in EOfAD-like and MPS IIIB mutants.

In the *ribosome* gene set, 48 leading edge genes were shared between MPS IIIB female (89%), EOfAD-like female (86%) and MPS IIIB male (64%) mutants (**Fig.5D**). This supports that gene set is altered in a similar way in these mutants. The magnitude of the fold change of the shared leading edge genes across all mutants was broadly larger in the MPS IIIB mutants compared to the EOfAD-like mutants (**Fig.S9**), consistent with MPS IIIB being an earlier-onset neurodegenerative disease.

We next sought to assess whether the changes to gene expression in the brains of MPS IIIB and EOfAD-like zebrafish were reflected at the protein level. To address this, we performed a proteomics experiment on 6-month-old wild type, EOfAD-like and MPS IIIB sibling brains. We detected up to 126,354 peptides per sample, corresponding to 9,942 unique proteins across the 30 samples.

We first performed PCA on the zebrafish brain proteomes to identify that the MPS IIIB proteomes generally formed a distinct cluster from the wild type and EOfAD-like genotypes (**Fig.6A-C**), similar to the RNA-seq data. We next tested which proteins were differentially expressed in each genotype and sex using moderated *t*-tests implemented in *limma* [78]. We found no statistical evidence supporting differentially expressed proteins in the EOfAD-like brains relative to their non-mutant siblings. However, the MPS IIIB females had 85 differentially expressed proteins, and the MPS IIIB males had 116 differentially expressed proteins relative to their wild type siblings (**Fig.6D,E,** full output in **Supplemental Data 4**). We then used STRING [79] to perform functional analysis of these differentially expressed proteins in MPS IIIB mutants. This revealed changes to lysosomal proteins and proteins involved in mTOR signalling in both sexes (**Fig.S10**).

**Fig. 6:**
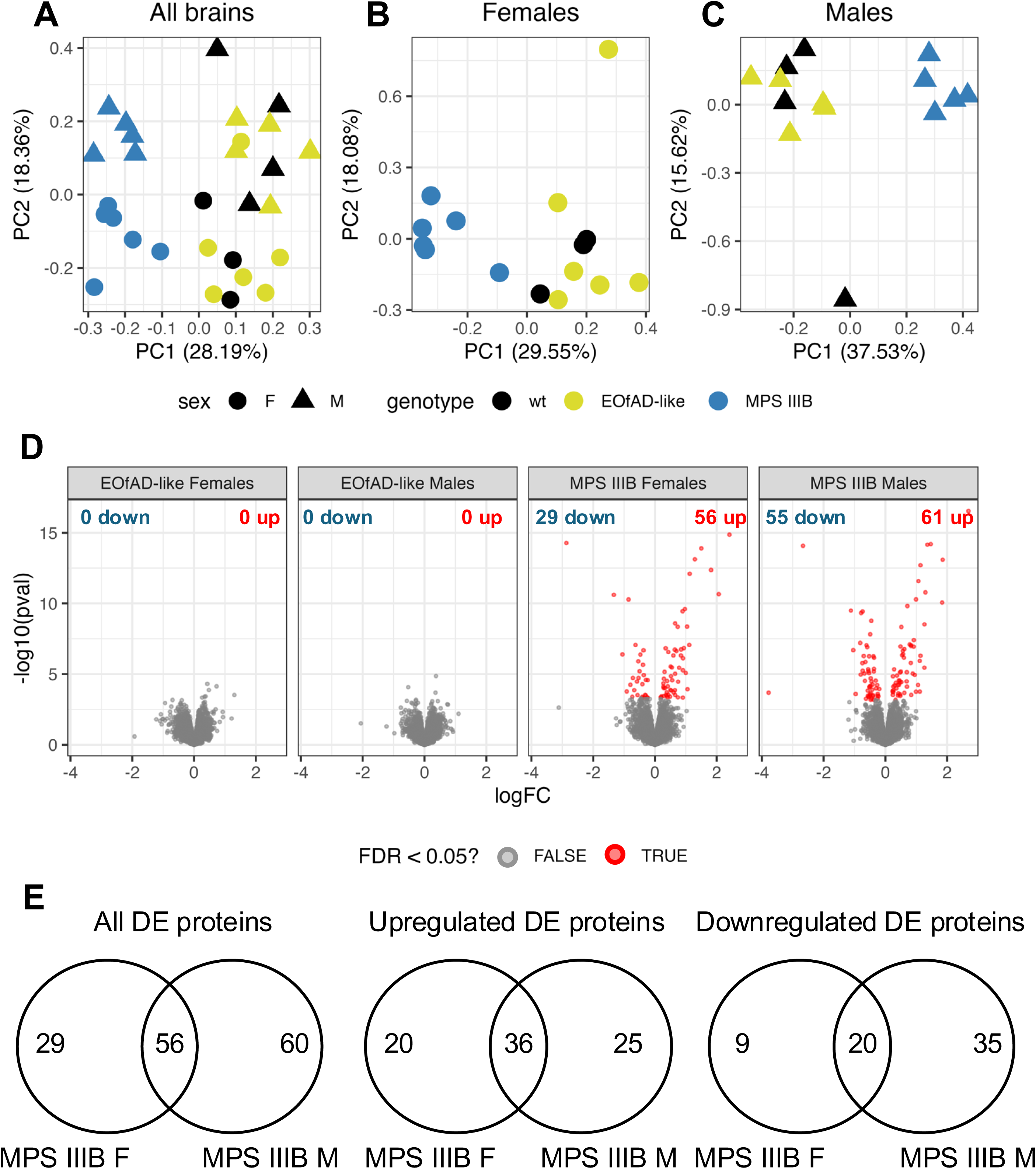
Differential protein expression analysis in zebrafish brains at 6 months of age. Principal component analysis (PCA) on protein abundance values after log2 transformation and quantile normalisation. Each point represents an adult zebrafish brain proteome sample in **A)** all samples, **B)** only female samples and **C)** only male samples that are coloured by genotype and shaped by sex (F: female, M: male, wt: wild type). The percent variations that principal component 1 (PC1) and 2 (PC2) explain in the dataset are indicated in parentheses in the axis labels. **D)** Volcano plots showing the log fold change (logFC) and negative log_10_ of the p-value of the proteins expressed in MPS IIIB and EOfAD-like zebrafish adult brains. Each point represents a protein which is coloured red if the gene was differentially expressed at an FDR-adjusted p-value of less that 0.05.

Since we did not identify significant altered proteins at the single protein level in EOfAD-like mutants, we subjected the proteomics data to the ensemble approach for gene set testing, where more subtle changes to protein expression can be detected. Using this approach, we detected statistical evidence for four gene sets to be altered in common in MPS IIIB and EOfAD-like female zebrafish, and two gene sets in males (**Fig.7A**, the full list of statistical significance of KEGG protein sets can be found in **Supplemental Data 3**). Out of the 3 gene sets significantly altered in both mutants in the transcriptome, only the lysosome and ribosome gene sets were also found to be significantly altered in both mutants in the proteome in at least one sex. We observed broad increased abundance of lysosomal proteins in MPS IIIB mutants (**Fig.7B**). Only 7 proteins were shared in the leading edge for the *lysosome* protein set across the MPS IIIB mutants and the female EOfAD-like mutants, and these proteins were altered in opposite directions (**Fig.S11A**). The *ribosome* protein set was broadly upregulated in both male mutants (**Fig.7C**), and the shared 25 proteins were altered in the same direction. Together, these results support that lysosomal proteins are altered in a different way in EOfAD-like and MPS IIIB mutants, and that ribosomal proteins are altered in a similar way.

**Fig. 7:**
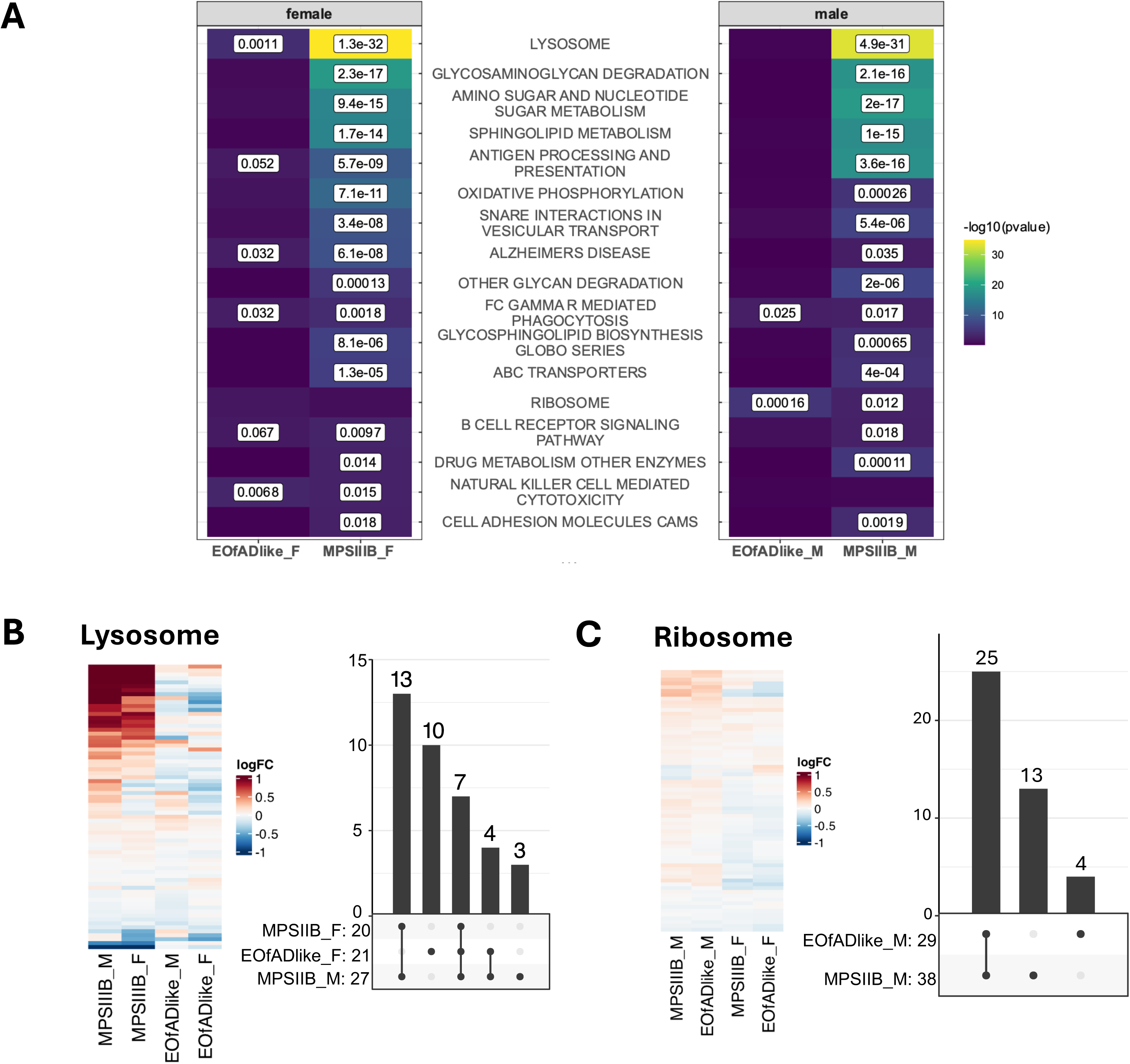
KEGG gene sets significantly altered in zebrafish brain proteomes at 6 months of age. **A)** Heatmap indicating those KEGG protein sets significantly altered in least two comparisons of mutant zebrafish to wild type. Cells in the heatmap are coloured by the log_10_ of the harmonic mean p-value (i.e. the brighter, more yellow cells are more statistically significant). The numbers indicate the FDR-adjusted p-values that were less than 0.1. **B)** Heatmaps indicating the similarities in the log fold change (logFC) of proteins in the KEGG protein sets for *lysosome* and **C)** *ribosome* in EOfAD-like and MPS IIIB mutant zebrafish brains. Upset plots display the overlap of leading edge proteins across the protein sets. The numbers next to each contrast represent the total number of leading-edge protein. The numbers above the bars indicate the number of protein shared between the protein sets in each contrast shown in the corresponding intersections.

The oxidative phosphorylation gene set was only significantly altered in MPS IIIB mutant proteomes (**Fig.7A**). However, we noted the particular downregulation of Atp6v0cb protein in MPS IIIB and female EOfAD-like mutants, an important component of the v-ATPase complex (**Fig.S12**) involved in lysosomal acidification [86]. We also observed significant changes to two pathways involved in innate immunity in EOfAD-like and MPS IIIB mutants that were not detected in the brain transcriptomes. The statistical significance of these two gene sets, *Fc gamma r mediated phagocytosis* and *natural killer cell mediated cytotoxicity* both are driven in part by the downregulation of the Rac2 protein (**Fig.S13**).

### Are changes in cell type proportions influencing the observed changes to gene expression?

In the analysis of bulk tissue, differences in the proportions of cell types between comparison groups may result in differences in transcript or protein abundances between those groups that are not related to changes in cellular functions. To assess whether this has occurred for the larval and adult brain tissues we analysed here, we utilised single cell RNA-seq datasets from 5 dpf zebrafish larvae [87] and from wild type (Tübingen) adult brain [88] to define gene sets consisting of markers of various cell types in zebrafish (we could not find a suitable dataset of 7 dpf zebrafish larvae in the literature). We then tested whether these marker gene sets were significantly enriched with up- or downregulated genes using *fry* [72] in both the transcriptomes and proteomes. No statistical evidence was found supporting changes to cell type gene sets in the EOfAD-like mutants at either age (**Fig.8A**, full output in **Supplemental Data 3**). However, significant changes to gene expression in marker genes of neural cell types were identified in 6-month-old MPS IIIB brains (**Fig.8A**). Notably, we observed a broad downregulation of genes expressed in oligodendrocytes (**Fig.8B**) and neural stem cells (**Fig.8C**) at the mRNA and protein level, suggesting that these cell types may show changes to their relative proportions. Additionally, inflammatory cell types (macrophages and microglia) showed changes to expression in MPS IIIB brains. However, a consistent direction of change was not observed (**Fig.8D,E**), supporting that these cell types may be altered, rather than changed in proportions. Collectively, these results indicate the presence of cell type-specific dysfunction in the brains of 6-month-old MPS IIIB zebrafish, potentially accompanied by alterations in the relative proportions of these cell types.

**Fig. 8.**
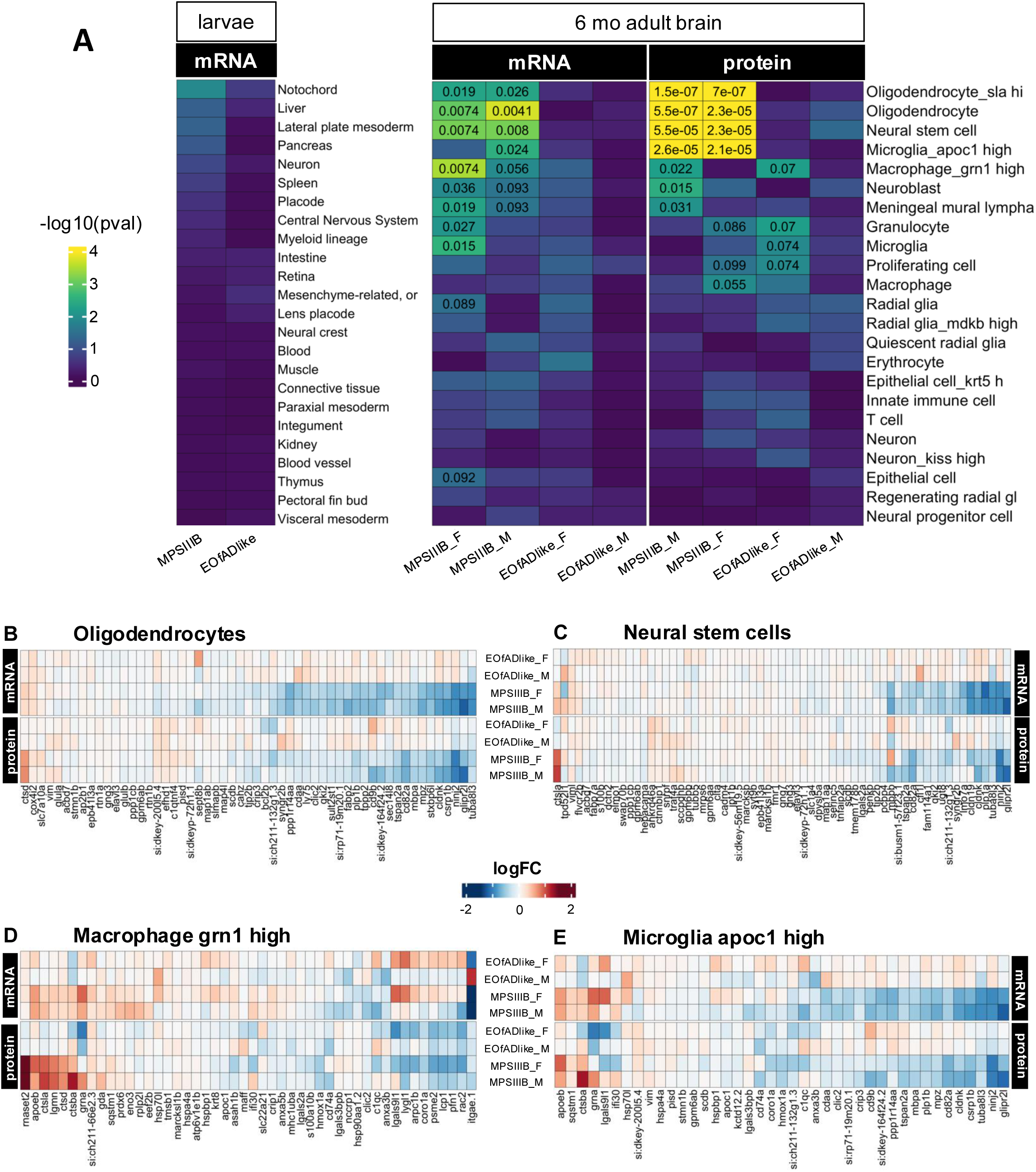
Possible changes to cell type proportions in MPS IIIB zebrafish brains. **A)** Heatmaps indicating the statistical significance of cell type marker gene sets in 7 days post fertilisation (dpf) larvae and 6 months of age (6m) adult brains. The cells in the heatmap are coloured by the log_10_ of the harmonic mean p-value (i.e. the brighter, more yellow cells are more statistically significant. The numbers indicate the FDR-adjusted p-values that were less than 0.1. **B)** Heatmap indicating the logFC values of the genes expressed in oligodendrocytes, **C)** neural stem cells, **D)** macrophages with high *grn1* expression and **E)** microglia with high *apoc1* expression.

### No observed effect on the expression of genes responsive to intracellular iron levels

Our previous analysis of the EOfAD-like mutation *psen1^Q96_K97del^* in 6-month-old female zebrafish brains [89] revealed significant changes in gene expression in genes regulated by intracellular iron levels. Specifically, the changes in expression occurred for genes containing iron-responsive elements (IREs) in the 3’ untranslated regions of their transcripts. However, we did not detect any statistical evidence for changes in expression of genes or proteins possessing IREs in either mutant at either age (**Fig. S14,** full output in **Supplemental Data 3**). Therefore, these datasets do not support that intracellular iron levels are abnormal in either of these mutants.

### Consistency with other changes in gene expression previously observed in EOfAD-like zebrafish

Our failure to observe changes in IRE gene sets in the 6-month-old EOfAD brains described here, led us to examine the consistency between all our analyses of the *psen1*^Q96_K97del^ mutation across independent experiments. We have previously analysed the effect of this mutation in n = 6 pools each of 40 larvae at 7 dpf [85], as well as in female 6-month-old zebrafish brains at n = 4 fish per genotype [54, 89]. To assess the overall similarity between samples across the experiments at each age, we performed PCA on the logCPM values of genes considered detectable in each experiment. This revealed that the samples separated based on their experiment of origin and not by genotype. This supports that the overall effect of genotype is very subtle in these samples, relative to the effect of experimental design (**Fig.9A,B**). We then applied our ensemble approach to gene set testing to each of the previous datasets to identify whether similar changes to the KEGG pathways could be identified. In 7 dpf larvae, this detected 85 significantly altered KEGG gene sets (**Fig.S15**), with the *ECM receptor interaction* gene set having the highest level of statistical significance (the 10 most significantly altered gene sets from the Dong et al. dataset [85] and their corresponding p-values from the current dataset are shown in **Fig.9C**). In the 6-month-old female adult brain dataset described in [54, 89], we detected 10 significantly altered KEGG gene sets, 6 of which were significantly altered in this experiment (all gene sets found to be significantly altered and their corresponding levels of statistical significance are shown in **Fig.9D**). Notably, there was a lack of correlations of the logFC values of the genes within these gene sets between the independent experiments (**Fig.9E-J**). Together, these results suggest that while extensive variation is apparent (particularly at the single gene level) common biological themes are still identified when the same mutation is analysed.

**Fig. 9.**
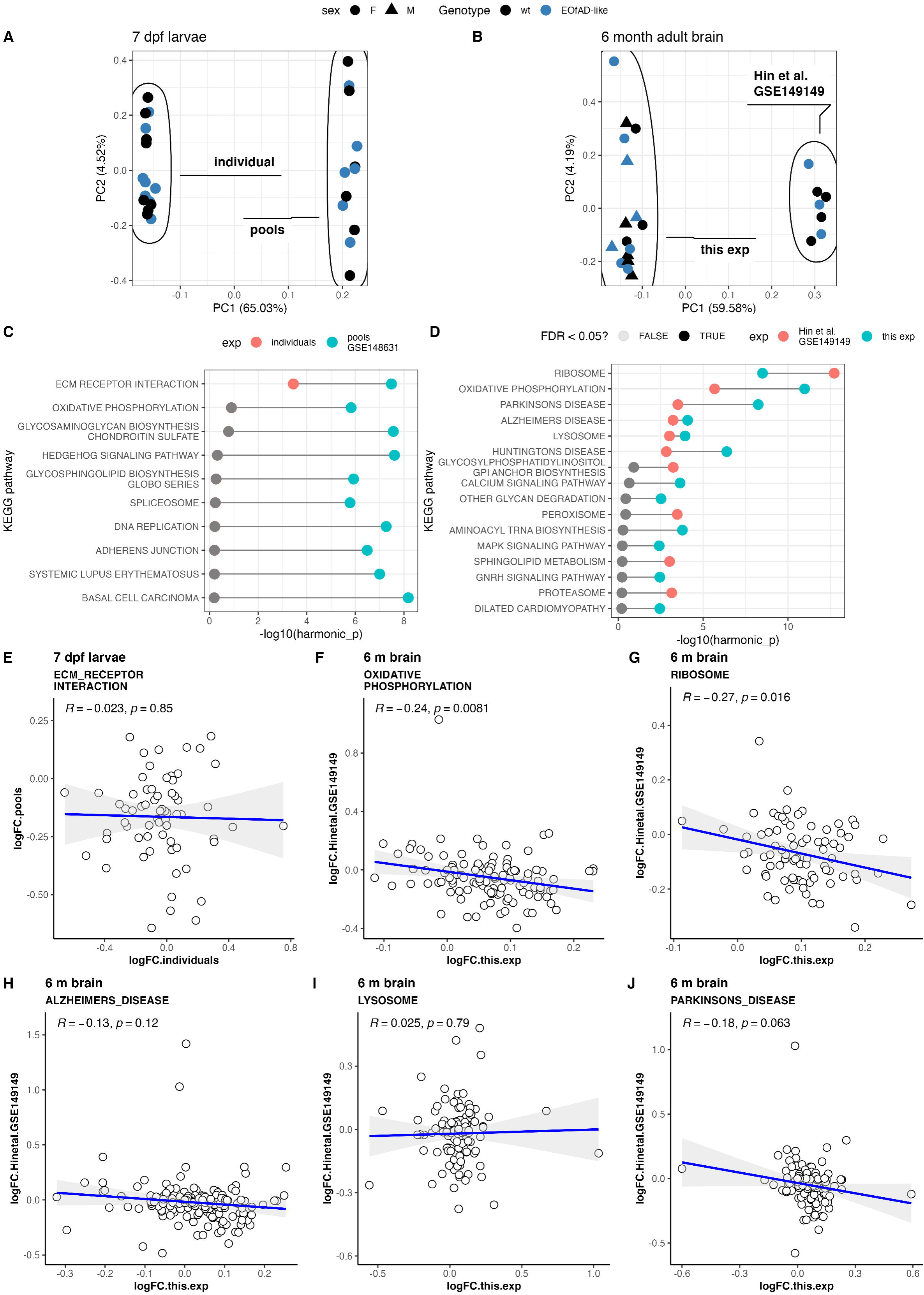
Consistent changes to KEGG pathways can be detected across independent experiments analysing EOfAD-like zebrafish. **A)** Principal component (PC) analysis of logCPM values in wild type (wt) and EOfAD-like zebrafish larvae at 7 days post fertilisation when analysing individuals (this experiment) or pools of 40 larvae [85]. **B)** PC analysis of logCPM values in 6-month-old wt (wild type) and EOfAD-like zebrafish brains in this experiment and from the data described in Hin et al. [89] **C)** Dumbbell plot showing the -log_10_ of the harmonic mean p-value of the 10 most significantly altered KEGG gene sets in 7 dpf individuals (red) or pooled (blue) EOfAD-like zebrafish larvae. **D)** Dumbbell plot showing the -log_10_ of the harmonic mean p-value of the KEGG gene sets significantly altered in the brains of 6-month-old EOfAD-like zebrafish in this experiment (red) or in the experiment described by Hin et al. [89] (blue). Points in **C)** and **D)** appear grey if they did not reach the threshold for being considered statistically significant at FDR < 0.05. **E)** Pearson correlation of the logFC values for genes in the KEGG gene set *ECM receptor interaction* in 7 dpf zebrafish larvae. **F)** Pearson correlation of the logFC values in 6-month-old EOfAD-like zebrafish brains for genes in the KEGG gene sets **F)** *oxidative phosphorylation*, **G)** *ribosome*, **H)** *Alzheimer’s disease*, **I)** *lysosome* and **J)** *Parkinson’s disease*.

## Discussion

In this study, we exploit the intra-family analysis strategy facilitated by zebrafish to provide, for the first time, a sensitive and direct comparison of genetic models of two inherited forms of early-onset dementia; EOfAD and MPS IIIB. We observed changes to gene expression in the extracellular matrix genes very early in these disease models, followed by changes to lysosomal and ribosomal gene and protein expression at later stages. Changes to oxidative phosphorylation gene expression was detected in 6 month old EOfAD-like and female MPS IIIB mutant brains, but this was only reflected in the MPS IIIB, and not EOfAD-like adult brain proteomes. We also identified changes to marker genes of specific neural cell types in MPS IIIB zebrafish brains, indicating these cell types may be particularly dysfunctional in this disease.

### Changes to ECM function could be an early, common disease mechanism in AD and MPS III

The extracellular matrix (ECM) is a complex network of proteins and carbohydrates that surrounds cells in multicellular organisms, providing mechanical support and playing a critical role in cellular functions such as tissue organisation and maintenance of tissue integrity. It consists primarily of proteins such as collagens and elastins, and glycoproteins such as laminins and fibronectin (reviewed in [90, 91]). Glycosaminoglycans incorporating heparan sulfate also form part of the ECM.

We observed subtle, yet statistically significant (**Fig.3A**), broad downregulation (**Fig.S7**) of the ECM genes in both MPS IIIB and EOfAD-like zebrafish larvae at 7 dpf. The ECM receptor interaction gene set was not significantly altered in 6 month old adult brain transcriptomes or proteomes, suggesting that ECM changes are restricted to larval zebrafish and/or are too subtle and not detectable when analysing the brains of young adults. The core pathology of MPS IIIB hinges on deficiency of NAGLU enzymatic activity, leading to lysosomal accumulation of the heparan sulfate degradation product. Extracellular accumulation of heparan sulfate has also been shown to occur in a *NAGLU* deficient cell line [92], and other extracellular substrates in other lysosomal storage disorders (reviewed in [93]), likely due to blockage of the endo-lysosomal system. Therefore, the broad downregulation of these ECM constituent genes in MPS IIIB zebrafish larvae suggests that the cells making up the bulk larval tissue are altering gene expression (presumably in an attempt to maintain homeostasis) in response to detection of excess ECM components, potentially due to impaired degradative capacity of lysosomes.

We observed a particular downregulation of the collagen genes in MPS IIIB larvae (**Fig.S7B**). Increased collagen IV (and laminin α-5) has been observed in MPS IIIA adenotonsillar tissue [94]. Interestingly, the formation of collagen I fibrils requires heparan sulfate [95]. Additionally, cathepsin K is a potent hydrolase of type I collagen [96], and the activity of this enzyme is inhibited by excess heparan sulfate [97]. Therefore, excess heparan sulfate leading to inhibition of cathepsin K may be interfering with collagen fibril formation in MPS IIIB larvae.

The role of the ECM in AD is not as clear. ECM proteins are upregulated in aged brains of 3xTg-AD mice [98], in hAPP-J20 mice [99] and in post-mortem AD brains [100, 101]. Increasing levels of extracellular matrix proteins are strongly associated with neuropathological traits and cognition in human AD brains relative to controls [102]. Additionally, they were found to be the earliest detectable biomarker in EOfAD patient cerebrospinal fluid proteomes, occurring 30 years before symptom onset [103].

The changes to ECM gene expression in EOfAD-like and MPS IIIB larvae are subtle. Nevertheless, they align with transcriptome analyses of several iPSC-derived neuron lines with *PSEN1* mutations [104–106], as well as in directly induced neurons (iNs) from AD [107], which more closely retain their epigenetic aging profiles than iPSC-derived neurons. Collagen type IV-encoding genes, which were downregulated in our zebrafish models, were found to be upregulated in MPS IIIB human fibroblasts relative to the other MPS III subtypes and MPS IV [108]. However, these collagen genes did not appear to be DE in comparison to a wild type control cell line [108]. This may be due to issues in the design of that transcriptome analysis such as low statistical power (one cell line per MPS disease) and an inappropriate control cell line likely preventing detection of significant changes to ECM-related gene expression. In contrast, iPSC-derived MPS IIIB neurons exhibited changes to the expression of ECM genes [109].

Together, our results and observations from the literature suggest that changes to the ECM play a role in the pathogenesis of MPS III and EOfAD. Our data indicate that these changes may occur very early in disease progression, may not be restricted to brain tissue, and potentially come about via similar mechanisms, as suggested by the similarity in changes to gene expression observed in zebrafish larvae (**Fig.3B**). Further investigation is required to confirm these mechanisms.

### Common changes to mitochondrial, lysosomal and ribosomal function occur in young adulthood

Zebrafish are recently sexually mature at 6 months of age, and we consider this equivalent to early adulthood in humans. For MPS IIIB, this is considered relatively late in disease progression as children typically succumb in their teenage years. This contrasts with EOfAD, where changes in cognition are not expected until the fourth decade of life or later. The dysregulation of genes associated with oxidative phosphorylation observed in the young adult brains of zebrafish models of EOfAD and MPS IIIB (**Fig.5A,B** and **Fig.S12**) suggests there may be a shared molecular pathway. However, the expression of proteins associated with oxidative phosphorylation was only observed to be altered in young adult MPS IIIB brain proteomes, not in EOfAD-like. Indeed, the magnitude of change at the gene expression level was minor in EOfAD-like zebrafish (**Fig.5B**) and so it is possible that these changes are not yet evident at the protein level.

Changes to the expression of the oxidative phosphorylation genes have been observed in young, pre-symptomatic animal models of AD. We previously analysed the effects of seven different AD-related mutations in 6-month-old zebrafish brains and found that these all resulted in changes to oxidative phosphorylation gene expression [48]. Mice engineered to express a humanised version of the human ε4 allele of the apolipoprotein E gene (*APOE4*, the strongest genetic risk factor for LOAD) also show changes to gene expression in the oxidative phosphorylation pathway at young ages [48, 110]. Impairment of this pathway has also been observed in diverse AD mouse models [111–114]. In humans, changes to the oxidative phosphorylation genes are found early in AD, both in blood [115] and in post-mortem brains [116]. Additionally, brain glucose metabolism (which generates metabolites to feed into oxidative phosphorylation) as measured by fluoro-D-glucose positron emission tomography (FDG-PET) consistently shows changes to glucose uptake during disease progression (reviewed in [117, 118]). Together, these observations from the literature imply that changes to mitochondrial function and oxidative phosphorylation occur early in AD. We suspect changes to oxidative phosphorylation proteins may become evident in later ages in EOfAD-like zebrafish brains.

The role of mitochondrial dysfunction in MPS III is not extensively characterised and how brain energy metabolism changes during disease progression in children with MPS III is poorly understood. Mitochondrial abnormalities and/or signs of oxidative stress (that can arise from dysfunctional mitochondrial respiration) have been observed in the brains of mouse models of MPS IIIA [119], IIIB [120], and IIIC [121–123]. Ultrastructural abnormalities [124] and signs of oxidative stress have also been observed in MPS IIIB human brain tissue [31]. A mouse model of MPS VII (lysosomal accumulation of several mucopolysaccharides including HS [125, 126]) showed a generally decreased abundance of the oxidative phosphorylation proteins which were not detected by microarray in age-matched samples of the same strain [127]. While the precise mechanism(s) of how and why mitochondria are affected throughout MPS III disease course is unclear, these may be consequences of defective autophagy/mitophagy (reviewed in [128]) caused by lysosome dysfunction.

Our analyses also detected significant changes to genes encoding lysosomal enzymes and proteins (**Fig.5**, **Fig.7**) in both EOfAD-like and MPS IIIB zebrafish brains. This gene set is broadly upregulated in both models at the mRNA level (**Fig.5C**). At the protein level, this gene set remains broadly upregulated in MPS IIIB brains, but the key genes driving the statistical significance of the gene set diverge in logFC in female EOfAD-like zebrafish (more than half of which are cathepsin proteases, **Fig.S11**). Lysosomes maintain an acidic lumen with pH values generally within a range of 4.5 – 5 [129] via a mechanism involving the v-ATPase complex (reviewed in [130, 131]). The genes encoding the components of this complex are broadly upregulated in both EOfAD-like and MPS IIIB zebrafish brains (in the final column cluster in **Fig.S12**). At the protein level, we observed downregulation of Atp6v0cb in MPS IIIB (only females) and EOfAD-like zebrafish brains. Atp6v0cb is a key component of the v-ATPase complex, forming the ring-like structure involved in the proton pumping mechanism (i.e. acidification) [132]. Aβ binds to the Atp6v0c subunit and this disrupts v-ATPase activity [133]. Knockdown of Atp6v0c causes accumulation of the C99 fragment of APP β-CTFs (i.e. the precursor to Aβ, [134]), which also inhibits lysosomal acidification [135]. The broad upregulation of these transcripts (**Fig.S12**), and the lower abundance of Atp6v0cb (which implies defective v-ATPase activity), may reflect a mechanism where the cells constituting these bulk brain RNA-seq samples are attempting to restore the normal pH of defective lysosomes to maintain cellular homeostasis. Lysosomal acidification defects have been detected in *PSEN1-*deficient and primary *PSEN1*-EOfAD fibroblasts [136], as well as several other models of AD mutations [135, 137–139]. Lysosomal pH was not detected as significantly altered in MPS IIIA mouse embryonic fibroblasts [140]. However, the existence of defective lysosomal acidification requires exploration in other MPS III subtypes, and in other models, particularly as it has been observed in several other lysosomal storage disorders [140–143] including MPS I [144].

Sufficient lysosomal acidification is required for the release of ferrous iron into the cytosol for absorption by mitochondria [145]. We did not observe significant changes to iron homeostasis-related genes in either mutant at either age (**Fig.S15**), despite previous detection of changes to iron transport gene expression in pools of EOfAD-like larvae at 7 dpf [85], changes in the *all 3’ IRE* gene set (*ire-3all*) in female EOfAD-like brains at 6 months of age [89], and in all IRE gene sets in MPS IIIA zebrafish brains at 3 months of age [47]. This was unexpected, and the reason(s) for this discrepancy is not fully understood (discussed later). However, there is evidence supporting iron dyshomeostasis in mild cognitive impairment (the precursor to AD) [146, 147], AD [148, 149], and in post-mortem AD brains [150, 151]. Iron homeostasis in MPS IIIB is far less studied. However, increased iron levels are observed in the cortex of a MPS IIIB mouse model in its symptomatic disease stages [152], as well as in a human MPS IIIB patient [153]. We suspect that changes to iron homeostasis genes in young adult zebrafish brains are very subtle and may be detected more consistently at later ages.

We observed broad, yet subtle, upregulation of ribosomal protein genes in EOfAD-like and MPS IIIB zebrafish brains (**Fig.5D**), and in male MPS IIIB and EOfAD-like proteomes. (**Fig.7A,C**). Upregulation of the ribosomal gene *RPLPA* was observed in a MPS IIIB patient-derived fibroblast line (and downregulation of the ribosomal genes *RPL10* and *RPL23* was observed in fibroblasts from other MPS III subtypes [108]). However, the functionality of ribosomes in MPS III is poorly understood.

Early in AD, protein synthesis rates by ribosomes appear to be decreased [154], possibly due to oxidation (by redox-active iron) of ribosomal proteins or RNA [155]. The observed, broad upregulation of the ribosomal gene set could reflect a homeostatic response to defects in protein synthesis in both EOfAD and MPS IIIB. Alternatively, we have speculated previously that it may be due to changes in mTOR signalling caused by changes in lysosomal function (discussed in [48]). This is supported by the dysregulation of mTOR proteins in MPS IIIB brain proteomes (**Fig.S10**).

### Cell type-specific changes to gene expression in MPS IIIB zebrafish brains

In MPS IIIB adult brains, we observed statistical evidence for changes to gene expression in four main cell types. The two most statistically significantly altered marker gene sets were those of oligodendrocytes and neural stem cells. These showed broad downregulation (**Fig.8B,C**). Significant reduction of the expression of the oligodendrocyte gene set was observed in the brains of our MPS IIIA, MPS IIIB and MPS IIIC zebrafish models at 3 months of age [47], as well as in a mouse model of MPS VII at approximately 5 months of age [156]. These gene expression changes are consistent with the observed demyelination in the cerebral cortex of MPS IIIB mice [157] and the corpus collosum of MPS IIIC mice [158] at 6 months of age. Intriguingly, mucopolysaccharides have been shown to inhibit oligodendrocyte maturation [159–161]. Together, our results along with previous literature of animal models of MPS diseases support that changes to oligodendrocyte differentiation and/or function occur due to the excess mucopolysaccharides in symptomatic MPS disease stages. In children with MPS III, loss of myelin [162, 163], and reduction of key oligodendrocyte proteins has been observed [158]. However, changes to oligodendrocyte function and myelination are not currently regarded as significant contributors to symptom manifestation.

We did not observe any evidence supporting oligodendrocyte dysfunction in EOfAD-like zebrafish at 7 dpf or 6 months of age. However, increasing evidence supports a role for oligodendrocytes in this disease. A brain imaging study of EOfAD patients revealed that white matter hyperintensities may be occurring as early as 22 years prior to symptom onset [164], which likely reflects demyelination and damage to the axons of neurons [165]. Intriguingly, expression of *PSEN1* in the human brain is highest in oligodendrocytes [166]. It is possible that oligodendrocytes in EOfAD-like zebrafish are affected, just not detectably at these ages.

We also observed significant changes to the expression of genes normally expressed in zebrafish macrophages and microglia (**Fig.8**), suggesting that inflammation is apparent in MPS IIIB brains. We also observed significant changes to inflammatory KEGG gene sets in the MPS IIIB brain proteomes: *Fc gamma r mediated phagocytosis* and *natural killer cell mediated cytotoxicity.* A clear up- or downregulation was not observed in the cell type gene sets, indicating they may not be under or over-represented in 6-month-old MPS IIB brains, rather their functionality may be altered. Intriguingly, microglia were observed to be the most affected cell type without any apparent changes in proportions in 8 month old MPS IIIA mouse brains in a single-nuclei RNA-seq analysis [167]. The statistical significance of the *Fc gamma r mediated phagocytosis* and *natural killer cell mediated cytotoxicity* gene sets in zebrafish MPS IIIB brain proteomes are, to an extent, driven by the downregulation of the protein Rac2. Rac2 is a small Rho GTPase expressed in leukocytes in adult zebrafish brains [168]. Rac2 is an essential co-factor in NADPH oxidase in leukocytes to form reactive oxygen species (ROS) inducing inflammation [169, 170]. While the role of Rac2 in zebrafish brains is not well characterised, its reduced abundance may imply that excessive ROS are present in these zebrafish brains (possibly generated by mitochondria via oxidative phosphorylation) potentially contributing to a neuroinflammatory phenotype.

### Consistent changes to gene expression can be detected at the pathway level across independent experiments

The changes to gene expression we observed in the EOfAD-like zebrafish in this experiment were consistent with our previous analyses [48, 54, 85, 89]. In each of those datasets, we observed significant changes to gene expression in pathways associated with the ECM in larvae, and in mitochondria, ribosomes and lysosomes in EOfAD-like adult brain. However, comparisons made at the single gene level did not reveal consistency (**Fig.9**). The differences at the single gene level may be attributed to technical factors inherent to conducting experiments independently, such as differences in sample preparation and sequencing platforms. Moreover, the inherent biological variability within the zebrafish models analysed at different times may lead to distinct outcomes when comparisons are not performed within a single experiment. Nevertheless, while single gene expression comparisons were inconsistent, the same pathways were generally identified to be significantly altered. This exemplifies the importance of interpreting results at the pathway or gene set level, where common pathways are identified, rather than at the single gene level when analysing transcriptome data.

## Limitations

In the work described in this paper, we have employed zebrafish as a model organism due to its unique advantages of being a commonly used vertebrate model species with high fecundity, genetic tractability, and fairly high genetic similarity to humans (reviewed in [171]). It is important to acknowledge that zebrafish may not fully recapitulate all aspects of human neurodegenerative diseases (particularly for AD), and the relative simplicity and small size of zebrafish brains may limit their equivalence to the human brain diseases we are attempting to model. Additionally, zebrafish are exceptionally regenerative, including the ability to regenerate brain tissue [172]. Therefore, zebrafish are not an ideal model to study the mechanisms of neurodegeneration specifically. Consequently, while our zebrafish models offer valuable insights into disease mechanisms preceding neurodegeneration, any findings should be interpreted with consideration of potential species-specific differences.

We observed broad downregulation of oligodendrocyte and neural stem cell marker genes in MPS IIIB zebrafish brains. This could be interpreted as indicating that the functions of these cell types are particularly disturbed by the MPS IIIB genotype. Alternatively, it could imply that there are significant changes in the proportions of these cell types constituting the MPS IIIB bulk brain transcriptome samples. In future work, we will dissect the changes to gene expression in MPS IIIB brains at the level of single cells to explore cell-type specific changes to gene expression, and whether there are changes to proportions of these cell types. Additionally, our findings will be verified with functional assays.

## Conclusion

Here, we provide the first direct comparison of zebrafish models of MPS IIIB and EOfAD, which lead to childhood and adult-onset dementia respectively. While the upstream genetic causes of these diseases are independent, our results suggest there may be some common disease mechanisms (summarised in **Table 2**). If this proves to be the case, common therapeutic strategies could be applied with benefit for people living with these devastating diseases.

**Table 2:** Summary of pathways altered in both MPS III and EOfAD-like zebrafish. . Only pathways altered in both mutants in larvae, and both the transcriptome and proteome in at least one sex of each mutant are listed.

| Pathway | Age/tissue | Transcriptome | Proteome | Comment |
| --- | --- | --- | --- | --- |
| ECM receptor interaction | 7 dpf | down | NA |  |
| Lysosome | 6 mo brain | upregulated | Up in MPS IIIB, mixed in EOfAD-like | not significant in EOfAD-like male proteomes |
| Ribosome | 6 mo brain | upregulated | upregulated | not significant in female brain proteomes |
| Oxidative phosphorylation | 6 mo brain | upregulated | upregulated | only significant in MPS IIIB brain proteomes |

## Declarations

### Ethics approval and consent to participate

All experiments involving zebrafish were conducted under the auspices of the Animal Ethics Committee of the University of Adelaide (approval numbers S-2021–067 and S-2021–041), and Institutional Biosafety Committee (PC2 NLRD 15037).

### Consent for publication

Not applicable.

### Availability of data and materials

The raw fastq files from these analyses have been deposited to the GEO database (https://www.ncbi.nlm.nih.gov/geo/) under the accession numbers GSE217196 (7 dpf larvae) and GSE242370 (adult brain). The data arising from our previous analyses of the EOfAD-like mutations can be obtained from the GEO database under accession numbers GSE148631 (pools of 7 dpf larvae) and GSE149149 (adult brain). The mass spectrometry proteomics data have been deposited to the ProteomeXchange Consortium via the PRIDE [173] partner repository with the dataset identifier PXD058264. All code to reproduce these analyses can be found at https://github.com/karissa-b/2021_MPSIIIBvQ96-RNAseq-7dpfLarve.

### Competing interests

The authors declare that they have no competing interests.

### Funding

This work was supported by a Postdoctoral Fellowship awarded to KB from Race Against Dementia and the Dementia Australia Research Foundation, and by funds from The Carthew Family Charity Trust to ML. KB is also supported by funds from Flinders University. ML and KH are academic employees of the University of Adelaide and Flinders University respectively.

### Authors’ contributions

KB and ML conceived the project. RB and MS generated the proteomics data. KB, RB and MS analysed the proteomics data. KB performed all other experiments and analyses described in this manuscript, prepared the figures, and wrote the first draft under the supervision of KH and ML. All authors read and contributed to the final manuscript.

## Acknowledgements

The authors would like to thank Lachlan Baer for his assistance with the snakemake pipeline. The authors acknowledge the South Australian Genomics Centre which provided RNA-seq library preparation and RNA sequencing services. The SAGC is supported by the National Collaborative Research Infrastructure Strategy (NCRIS) via BioPlatforms Australia and by the SAGC partner institutes. This work was supported with supercomputing resources provided by the Phoenix HPC service at the University of Adelaide.

## Notes

### Competing Interest Statement

The authors have declared no competing interest.

### Summary of Updates

added proteomics and changes to transcriptome analysis to incorporate sex differences

## References

1. Rizzi L, Rosset I, Roriz-Cruz M: Global epidemiology of dementia: Alzheimer’s and vascular types. Biomed Res Int 2014, 2014:908915.

2. Alzheimer A: Uber eigenartige Erkrankung der Hirnrinde. All Z Psychiatr 1907, 64:146–148.

3. Du X, Wang X, Geng M: Alzheimer’s disease hypothesis and related therapies. Transl Neurodegener 2018, 7:2.

4. Kepp KP, Robakis NK, Høilund-Carlsen PF, Sensi SL, Vissel B: The amyloid cascade hypothesis: an updated critical review. Brain 2023:awad159.

5. Sherrington R, Rogaev EI, Liang Y, Rogaeva EA, Levesque G, Ikeda M, Chi H, Lin C, Li G, Holman K et al: Cloning of a gene bearing missense mutations in early-onset familial Alzheimer’s disease. Nature 1995, 375(6534):754-760.

6. Rogaev EI, Sherrington R, Rogaeva EA, Levesque G, Ikeda M, Liang Y, Chi H, Lin C, Holman K, Tsuda T et al: Familial Alzheimer’s disease in kindreds with missense mutations in a gene on chromosome 1 related to the Alzheimer’s disease type 3 gene. Nature 1995, 376(6543):775–778.

7. Levy-Lahad E, Wasco W, Poorkaj P, Romano DM, Oshima J, Pettingell WH, Yu C-e, Jondro PD, Schmidt SD, Wang K et al: Candidate Gene for the Chromosome 1 Familial Alzheimer’s Disease Locus. Science 1995, 269(5226):973–977.

8. Goate A, Chartier-Harlin M-C, Mullan M, Brown J, Crawford F, Fidani L, Giuffra L, Haynes A, Irving N, James L et al: Segregation of a missense mutation in the amyloid precursor protein gene with familial Alzheimer’s disease. Nature 1991, 349(6311):704–706.

9. Pottier C, Hannequin D, Coutant S, Rovelet-Lecrux A, Wallon D, Rousseau S, Legallic S, Paquet C, Bombois S, Pariente J et al: High frequency of potentially pathogenic SORL1 mutations in autosomal dominant early-onset Alzheimer disease. Molecular Psychiatry 2012, 17(9):875–879.

10. Nicolas G, Charbonnier C, Wallon D, Quenez O, Bellenguez C, Grenier-Boley B, Rousseau S, Richard AC, Rovelet-Lecrux A, Le Guennec K et al: SORL1 rare variants: a major risk factor for familial early-onset Alzheimer’s disease. Molecular Psychiatry 2016, 21(6):831–836.

11. Pasternak SH, Bagshaw RD, Guiral M, Zhang S, Ackerley CA, Pak BJ, Callahan JW, Mahuran DJ: Presenilin-1, nicastrin, amyloid precursor protein, and gamma-secretase activity are co-localized in the lysosomal membrane. J Biol Chem 2003, 278(29):26687–26694.

12. Kawai M, Cras P, Richey P, Tabaton M, Lowery DE, Gonzalez-DeWhitt PA, Greenberg BD, Gambetti P, Perry G: Subcellular localization of amyloid precursor protein in senile plaques of Alzheimer’s disease. The American journal of pathology 1992, 140(4):947–958.

13. Mishra S, Knupp A, Szabo MP, Williams CA, Kinoshita C, Hailey DW, Wang Y, Andersen OM, Young JE: The Alzheimer’s gene SORL1 is a regulator of endosomal traffic and recycling in human neurons. Cell Mol Life Sci 2022, 79(3):162.

14. Fedeli C, Filadi R, Rossi A, Mammucari C, Pizzo P: PSEN2 (presenilin 2) mutants linked to familial Alzheimer disease impair autophagy by altering Ca(2+) homeostasis. Autophagy 2019, 15(12):2044–2062.

15. Gao S, Casey AE, Sargeant TJ, Mäkinen V-P: Genetic variation within endolysosomal system is associated with late-onset Alzheimer’s disease. Brain 2018, 141(9):2711–2720.

16. Cataldo AM, Peterhoff CM, Troncoso JC, Gomez-Isla T, Hyman BT, Nixon RA: Endocytic pathway abnormalities precede amyloid beta deposition in sporadic Alzheimer’s disease and Down syndrome: differential effects of APOE genotype and presenilin mutations. The American journal of pathology 2000, 157(1):277–286.

17. Serrano-Pozo A, Frosch MP, Masliah E, Hyman BT: Neuropathological alterations in Alzheimer disease. Cold Spring Harb Perspect Med 2011, 1(1):a006189.

18. McKean NE, Handley RR, Snell RG: A Review of the Current Mammalian Models of Alzheimer’s Disease and Challenges That Need to Be Overcome. Int J Mol Sci 2021, 22(23).

19. Drummond E, Wisniewski T: Alzheimer’s disease: experimental models and reality. Acta Neuropathol 2017, 133(2):155–175.

20. Hargis KE, Blalock EM: Transcriptional signatures of brain aging and Alzheimer’s disease: What are our rodent models telling us? Behavioural Brain Research 2017, 322:311–328.

21. Mullane K, Williams M: Preclinical Models of Alzheimer’s Disease: Relevance and Translational Validity. Current Protocols in Pharmacology 2019, 84(1):e57.

22. Lardelli M: An Alternative View of Familial Alzheimer’s Disease Genetics. Journal of Alzheimer’s Disease 2023, Preprint:1-27.

23. Zelei T, Csetneki K, Vokó Z, Siffel C: Epidemiology of Sanfilippo syndrome: results of a systematic literature review. Orphanet journal of rare diseases 2018, 13(1):53.

24. Scott HS, Blanch L, Guo X-H, Freeman C, Orsborn A, Baker E, Sutherland GR, Morris CP, Hopwood JJ: Cloning of the sulphamidase gene and identification of mutations in Sanfilippo A syndrome. Nature genetics 1995, 11(4):465–467.

25. Zhao HG, Li HH, Bach G, Schmidtchen A, Neufeld EF: The molecular basis of Sanfilippo syndrome type B. Proceedings of the National Academy of Sciences 1996, 93(12):6101–6105.

26. Klein U, Kresse H, von Figura K: Sanfilippo syndrome type C: deficiency of acetyl-CoA:alpha-glucosaminide N-acetyltransferase in skin fibroblasts. Proceedings of the National Academy of Sciences 1978, 75(10):5185–5189.

27. Kresse H, Paschke E, von Figura K, Gilberg W, Fuchs W: Sanfilippo disease type D: deficiency of N-acetylglucosamine-6-sulfate sulfatase required for heparan sulfate degradation. Proceedings of the National Academy of Sciences 1980, 77(11):6822–6826.

28. Kowalewski B, Lamanna WC, Lawrence R, Damme M, Stroobants S, Padva M, Kalus I, Frese M-A, Lübke T, Lüllmann-Rauch R et al: Arylsulfatase G inactivation causes loss of heparan sulfate 3-O-sulfatase activity and mucopolysaccharidosis in mice. Proceedings of the National Academy of Sciences 2012, 109(26):10310–10315.

29. Wiśniewska K, Wolski J, Żabińska M, Szulc A, Gaffke L, Pierzynowska K, Węgrzyn G: Mucopolysaccharidosis Type IIIE: A Real Human Disease or a Diagnostic Pitfall? Diagnostics 2024, 14(16):1734.

30. Villani GRD, Domenico CD, Musella A, Cecere F, Napoli DD, Natale PD: Mucopolysaccharidosis IIIB: Oxidative damage and cytotoxic cell involvement in the neuronal pathogenesis. Brain Research 2009, 1279:99–108.

31. Hamano K, Hayashi M, Shioda K, Fukatsu R, Mizutani S: Mechanisms of neurodegeneration in mucopolysaccharidoses II and IIIB: analysis of human brain tissue. Acta Neuropathol 2008, 115(5):547–559.

32. Jones MZ, Alroy J, Rutledge JC, Taylor JW, Alvord EC, Jr., Toone J, Applegarth D, Hopwood JJ, Skutelsky E, Ianelli C et al: Human Mucopolysaccharidosis IIID: Clinical, Biochemical, Morphological and Immunohistochemical Characteristics. Journal of Neuropathology & Experimental Neurology 1997, 56(10):1158–1167.

33. Valle DAd, Santos MLSF, Telles BA, Cordeiro ML: Neurological, neurobehavioral, and radiological alterations in patients with mucopolysaccharidosis III (Sanfilippo’s syndrome) in Brazil. Frontiers in Neurology 2022, 13.

34. Ohmi K, Kudo LC, Ryazantsev S, Zhao H-Z, Karsten SL, Neufeld EF: Sanfilippo syndrome type B, a lysosomal storage disease, is also a tauopathy. Proceedings of the National Academy of Sciences 2009, 106(20):8332–8337.

35. Ginsberg SD, Galvin JE, Lee VMY, Rorke LB, et al.: Accumulation of intracellular amyloid-beta peptide (Abeta 1-40) in mucopolysaccharidosis brains. Journal of Neuropathology and Experimental Neurology 1999, 58(8):815–824.

36. Beard H, Hassiotis S, Gai W-P, Parkinson-Lawrence E, Hopwood JJ, Hemsley KM: Axonal dystrophy in the brain of mice with Sanfilippo syndrome. Experimental Neurology 2017, 295:243–255.

37. Viana GM, Priestman DA, Platt FM, Khan S, Tomatsu S, Pshezhetsky AV: Brain Pathology in Mucopolysaccharidoses (MPS) Patients with Neurological Forms. J Clin Med 2020, 9(2).

38. Howe K, Clark MD, Torroja CF, Torrance J, Berthelot C, Muffato M, Collins JE, Humphray S, McLaren K, Matthews L et al: The zebrafish reference genome sequence and its relationship to the human genome. Nature 2013, 496(7446):498–503.

39. Herrero J, Muffato M, Beal K, Fitzgerald S, Gordon L, Pignatelli M, Vilella AJ, Searle SM, Amode R, Brent S et al: Ensembl comparative genomics resources. Database (Oxford) 2016, 2016.

40. Haynes EM, Ulland TK, Eliceiri KW: A Model of Discovery: The Role of Imaging Established and Emerging Non-mammalian Models in Neuroscience. Frontiers in Molecular Neuroscience 2022, 15.

41. Liu Y: Zebrafish as a Model Organism for Studying Pathologic Mechanisms of Neurodegenerative Diseases and other Neural Disorders. Cellular and Molecular Neurobiology 2023.

42. Barthelson K, Dong Y, Newman M, Lardelli M: PRESENILIN 1 mutations causing early-onset familial Alzheimer’s disease or familial acne inversa differ in their effects on genes facilitating energy metabolism and signal transduction. Journal of Alzheimer’s Disease 2021, 82(1):327–347.

43. Barthelson K, Pederson SM, Newman M, Jiang H, Lardelli M: In-frame and frameshift mutations in zebrafish Presenilin 2 affect different cellular functions in young adult brains. Journal of Alzheimer’s Disease Reports 2021, 5(1):395–404.

44. Barthelson K, Pederson SM, Newman M, Lardelli M: Brain transcriptome analysis reveals subtle effects on mitochondrial function and iron homeostasis of mutations in the SORL1 gene implicated in early onset familial Alzheimer’s disease. Molecular Brain 2020, 13(1):142.

45. Barthelson K, Pederson SM, Newman M, Lardelli M: Brain transcriptome analysis of a protein-truncating mutation in sortilin-related receptor 1 associated with early-onset familial Alzheimer’s disease indicates early effects on mitochondrial and ribosome function. Journal of Alzheimer’s Disease 2021, 79(3):1105–1119.

46. Jiang H, Pederson SM, Newman M, Dong Y, Barthelson K, Lardelli M: Transcriptome analysis indicates dominant effects on ribosome and mitochondrial function of a premature termination codon mutation in the zebrafish gene psen2. Plos one 2020, 15(7):e0232559.

47. Gerken E, Ahmad S, Barthelson K, Lardelli M: Zebrafish models of Mucopolysaccharidosis types IIIA, B, and C show hyperactivity and changes in oligodendrocyte state. bioRxiv 2023:2023.2008.2002.550904.

48. Barthelson K, Newman M, Lardelli M: Brain transcriptomes of zebrafish and mouse Alzheimer’s disease knock-in models imply early disrupted energy metabolism. Disease Models & Mechanisms 2022, 15(1):dmm049187.

49. Lazic SE, Essioux L: Improving basic and translational science by accounting for litter-to-litter variation in animal models. BMC Neuroscience 2013, 14(1):37.

50. Armant O, Gourain V, Etard C, Strähle U: Whole transcriptome data analysis of zebrafish mutants affecting muscle development. Data in Brief 2016, 8:61–68.

51. Hu W, Yang S, Shimada Y, Münch M, Marín-Juez R, Meijer AH, Spaink HP: Infection and RNA-seq analysis of a zebrafish tlr2 mutant shows a broad function of this toll-like receptor in transcriptional and metabolic control and defense to Mycobacterium marinum infection. BMC Genomics 2019, 20(1):878.

52. Livne H, Avital T, Ruppo S, Harazi A, Mitrani-Rosenbaum S, Daya A: Generation and characterization of a novel gne Knockout Model in Zebrafish. Frontiers in Cell and Developmental Biology 2022, 10.

53. Weinschutz Mendes H, Neelakantan U, Liu Y, Fitzpatrick SE, Chen T, Wu W, Pruitt A, Jin DS, Jamadagni P, Carlson M et al: High-throughput functional analysis of autism genes in zebrafish identifies convergence in dopaminergic and neuroimmune pathways. Cell reports 2023, 42(3).

54. Newman M, Hin N, Pederson S, Lardelli M: Brain transcriptome analysis of a familial Alzheimer’s disease-like mutation in the zebrafish presenilin 1 gene implies effects on energy production. Mol Brain 2019, 12(1):43.

55. Westerfield M: The zebrafish book: a guide for the laboratory use of zebrafish. http://zfin org/zf_info/zfbook/zfbk html 2000.

56. Allen AG, Barthelson K, Lardelli M: pHAPE: a plasmid for production of DNA size marker ladders for gel electrophoresis. Biology Methods and Protocols 2023, 8(1):bpad015.

57. Mölder F, Jablonski KP, Letcher B: Sustainable data analysis with Snakemake.[version 2; peer review: 2 approved]. F1000Research 10: 33. Crossref, PubMed 2021.

58. Chen S, Zhou Y, Chen Y, Gu J: fastp: an ultra-fast all-in-one FASTQ preprocessor. Bioinformatics 2018, 34(17):i884–i890.

59. Dobin A, Davis CA, Schlesinger F, Drenkow J, Zaleski C, Jha S, Batut P, Chaisson M, Gingeras TR: STAR: ultrafast universal RNA-seq aligner. Bioinformatics 2013, 29(1):15–21.

60. Li H, Handsaker B, Wysoker A, Fennell T, Ruan J, Homer N, Marth G, Abecasis G, Durbin R, Genome Project Data Processing S: The Sequence Alignment/Map format and SAMtools. Bioinformatics 2009, 25(16):2078–2079.

61. Smith TS, Heger A, Sudbery I: UMI-tools: Modelling sequencing errors in Unique Molecular Identifiers to improve quantification accuracy. Genome Research 2017.

62. Liao Y, Smyth GK, Shi W: featureCounts: an efficient general purpose program for assigning sequence reads to genomic features. Bioinformatics 2014, 30(7):923–930.

63. Andrews S: FastQC: a quality control tool for high throughput sequence data. In.: Babraham Bioinformatics, Babraham Institute, Cambridge, United Kingdom; 2010.

64. Ward CM, To TH, Pederson SM: ngsReports: a Bioconductor package for managing FastQC reports and other NGS related log files. Bioinformatics 2020, 36(8):2587–2588.

65. Team RC: R: A Language and Environment for Statistical Computing. In. Edited by Computing RFfS. Vienna, Austria; 2020.

66. Robinson MD, McCarthy DJ, Smyth GK: edgeR: a Bioconductor package for differential expression analysis of digital gene expression data. Bioinformatics 2010, 26(1):139–140.

67. Hansen KD, Irizarry RA, Wu Z: Removing technical variability in RNA-seq data using conditional quantile normalization. *Biostatistics (Oxford*, England) 2012, 13(2):204–216.

68. Ashburner M, Ball CA, Blake JA, Botstein D, Butler H, Cherry JM, Davis AP, Dolinski K, Dwight SS, Eppig JT et al: Gene ontology: tool for the unification of biology. The Gene Ontology Consortium. Nature genetics 2000, 25(1):25–29.

69. Consortium GO: The Gene Ontology resource: enriching a GOld mine. Nucleic Acids Res 2021, 49(D1):D325–d334.

70. Young MD, Wakefield MJ, Smyth GK, Oshlack A: Gene ontology analysis for RNA-seq: accounting for selection bias. Genome Biology 2010, 11(2):R14.

71. Kanehisa M, Goto S: KEGG: Kyoto Encyclopedia of Genes and Genomes. Nucleic Acids Research 2000, 28(1):27–30.

72. Wu D, Lim E, Vaillant F, Asselin-Labat M-L, Visvader JE, Smyth GK: ROAST: rotation gene set tests for complex microarray experiments. Bioinformatics 2010, 26(17):2176–2182.

73. Wu D, Smyth GK: Camera: a competitive gene set test accounting for inter-gene correlation. Nucleic Acids Research 2012, 40(17):e133–e133.

74. Korotkevich G, Sukhov V, Budin N, Shpak B, Artyomov MN, Sergushichev A: Fast gene set enrichment analysis. bioRxiv 2021:060012.

75. Subramanian A, Tamayo P, Mootha VK, Mukherjee S, Ebert BL, Gillette MA, Paulovich A, Pomeroy SL, Golub TR, Lander ES, Mesirov JP: Gene set enrichment analysis: A knowledge-based approach for interpreting genome-wide expression profiles. Proceedings of the National Academy of Sciences 2005, 102(43):15545–15550.

76. Wilson DJ: The harmonic mean p-value for combining dependent tests. Proceedings of the National Academy of Sciences 2019, 116(4):1195–1200.

77. Bi R, Liu P: Sample size calculation while controlling false discovery rate for differential expression analysis with RNA-sequencing experiments. BMC bioinformatics 2016, 17:146.

78. Ritchie ME, Phipson B, Wu D, Hu Y, Law CW, Shi W, Smyth GK: limma powers differential expression analyses for RNA-sequencing and microarray studies. Nucleic Acids Res 2015, 43(7):e47.

79. Szklarczyk D, Kirsch R, Koutrouli M, Nastou K, Mehryary F, Hachilif R, Gable AL, Fang T, Doncheva NT, Pyysalo S et al: The STRING database in 2023: protein-protein association networks and functional enrichment analyses for any sequenced genome of interest. Nucleic Acids Res 2023, 51(D1):D638–d646.

80. Wickham H: ggplot2: Elegant Graphics for Data Analysis: Springer-Verlag New York; 2016.

81. Mangiola S, Papenfuss AT: tidyHeatmap: an R package for modular heatmap production based on tidy principles. Journal of Open Source Software 2020, 5(52):2472.

82. Luo W, Brouwer C: Pathview: an R/Bioconductor package for pathway-based data integration and visualization. Bioinformatics 2013, 29(14):1830–1831.

83. Conway JR, Lex A, Gehlenborg N: UpSetR: an R package for the visualization of intersecting sets and their properties. Bioinformatics 2017, 33(18):2938–2940.

84. Fedele AO: Sanfilippo syndrome: causes, consequences, and treatments. The application of clinical genetics 2015, 8:269–281.

85. Dong Y, Newman M, Pederson SM, Barthelson K, Hin N, Lardelli M: Transcriptome analyses of 7-day-old zebrafish larvae possessing a familial Alzheimer’s disease-like mutation in psen1 indicate effects on oxidagtive phosphorylation, ECM and MCM functions, and iron homeostasis. BMC Genomics 2021, 22(1):211.

86. Forgac M: Vacuolar ATPases: rotary proton pumps in physiology and pathophysiology. Nature Reviews Molecular Cell Biology 2007, 8(11):917–929.

87. Farnsworth DR, Saunders LM, Miller AC: A single-cell transcriptome atlas for zebrafish development. Developmental Biology 2020, 459(2):100–108.

88. Jiang M, Xiao Y, E W, Ma L, Wang J, Chen H, Gao C, Liao Y, Guo Q, Peng J et al: Characterization of the Zebrafish Cell Landscape at Single-Cell Resolution. Frontiers in Cell and Developmental Biology 2021, 9.

89. Hin N, Newman M, Pederson S, Lardelli M: Iron responsive element-mediated responses to iron dyshomeostasis in Alzheimer’s disease. Journal of Alzheimer’s Disease 2021, 84(4):1597–1630.

90. Karamanos NK, Theocharis AD, Piperigkou Z, Manou D, Passi A, Skandalis SS, Vynios DH, Orian-Rousseau V, Ricard-Blum S, Schmelzer CEH et al: A guide to the composition and functions of the extracellular matrix. The FEBS Journal 2021, 288(24):6850–6912.

91. Theocharis AD, Skandalis SS, Gialeli C, Karamanos NK: Extracellular matrix structure. Advanced Drug Delivery Reviews 2016, 97:4–27.

92. Roy E, Bruyère J, Flamant P, Bigou S, Ausseil J, Vitry S, Heard JM: GM130 gain-of-function induces cell pathology in a model of lysosomal storage disease. Human Molecular Genetics 2012, 21(7):1481–1495.

93. Batzios SP, Zafeiriou DI, Papakonstantinou E: Extracellular matrix components: An intricate network of possible biomarkers for lysosomal storage disorders? FEBS Letters 2013, 587(8):1258–1267.

94. Pal AR, Mercer J, Jones SA, Bruce IA, Bigger BW: Substrate accumulation and extracellular matrix remodelling promote persistent upper airway disease in mucopolysaccharidosis patients on enzyme replacement therapy. PLOS ONE 2018, 13(9):e0203216.

95. Hill KE, Lovett BM, Schwarzbauer JE: Heparan sulfate is necessary for the early formation of nascent fibronectin and collagen I fibrils at matrix assembly sites. Journal of Biological Chemistry 2022, 298(1):101479.

96. Brömme D, Okamoto K, Wang BB, Biroc S: Human cathepsin O2, a matrix protein-degrading cysteine protease expressed in osteoclasts. Functional expression of human cathepsin O2 in Spodoptera frugiperda and characterization of the enzyme. J Biol Chem 1996, 271(4):2126–2132.

97. Zhang X, Luo Y, Hao H, Krahn JM, Su G, Dutcher R, Xu Y, Liu J, Pedersen LC, Xu D: Heparan sulfate selectively inhibits the collagenase activity of cathepsin K. Matrix Biol 2024, 129:15–28.

98. Bourasset F, Ouellet M, Tremblay C, Julien C, Do TM, Oddo S, LaFerla F, Calon F: Reduction of the cerebrovascular volume in a transgenic mouse model of Alzheimer’s disease. Neuropharmacology 2009, 56(4):808–813.

99. Cheng JS, Dubal DB, Kim DH, Legleiter J, Cheng IH, Yu G-Q, Tesseur I, Wyss-Coray T, Bonaldo P, Mucke L: Collagen VI protects neurons against Aβ toxicity. Nature Neuroscience 2009, 12(2):119–121.

100. Kalaria RN, Pax AB: Increased collagen content of cerebral microvessels in Alzheimer’s disease. Brain Res 1995, 705(1-2):349–352.

101. Lepelletier FX, Mann DMA, Robinson AC, Pinteaux E, Boutin H: Early changes in extracellular matrix in Alzheimer’s disease. Neuropathology and Applied Neurobiology 2017, 43(2):167–182.

102. Johnson ECB, Carter EK, Dammer EB, Duong DM, Gerasimov ES, Liu Y, Liu J, Betarbet R, Ping L, Yin L et al: Large-scale deep multi-layer analysis of Alzheimer’s disease brain reveals strong proteomic disease-related changes not observed at the RNA level. Nat Neurosci 2022, 25(2):213–225.

103. Johnson ECB, Bian S, Haque RU, Carter EK, Watson CM, Gordon BA, Ping L, Duong DM, Epstein MP, McDade E et al: Cerebrospinal fluid proteomics define the natural history of autosomal dominant Alzheimer’s disease. Nature Medicine 2023, 29(8):1979–1988.

104. Kwart D, Gregg A, Scheckel C, Murphy EA, Paquet D, Duffield M, Fak J, Olsen O, Darnell RB, Tessier-Lavigne M: A Large Panel of Isogenic APP and PSEN1 Mutant Human iPSC Neurons Reveals Shared Endosomal Abnormalities Mediated by APP β-CTFs, Not Aβ. Neuron 2019, 104(2):256–270.e255.

105. Corsi GI, Gadekar VP, Haukedal H, Doncheva NT, Anthon C, Ambardar S, Palakodeti D, Hyttel P, Freude K, Seemann SE, Gorodkin J: The transcriptomic landscape of neurons carrying PSEN1 mutations reveals changes in extracellular matrix components and non-coding gene expression. Neurobiology of Disease 2023, 178:105980.

106. Caldwell AB, Liu Q, Zhang C, Schroth GP, Galasko DR, Rynearson KD, Tanzi RE, Yuan SH, Wagner SL, Subramaniam S: Endotype reversal as a novel strategy for screening drugs targeting familial Alzheimer’s disease. Alzheimer’s & Dementia 2022, 18(11):2117–2130.

107. Mertens J, Herdy JR, Traxler L, Schafer ST, Schlachetzki JCM, Böhnke L, Reid DA, Lee H, Zangwill D, Fernandes DP et al: Age-dependent instability of mature neuronal fate in induced neurons from Alzheimer&#x2019;s patients. Cell Stem Cell 2021, 28(9):1533–1548.e1536.

108. Wiśniewska K, Gaffke L, Krzelowska K, Węgrzyn G, Pierzynowska K: Differences in gene expression patterns, revealed by RNA-seq analysis, between various Sanfilippo and Morquio disease subtypes. Gene 2022, 812:146090.

109. Lemonnier T, Blanchard S, Toli D, Roy E, Bigou S, Froissart R, Rouvet I, Vitry S, Heard JM, Bohl D: Modeling neuronal defects associated with a lysosomal disorder using patient-derived induced pluripotent stem cells. Human Molecular Genetics 2011, 20(18):3653–3666.

110. Lee S, Devanney NA, Golden LR, Smith CT, Schwartz JL, Walsh AE, Clarke HA, Goulding DS, Allenger EJ, Morillo-Segovia G et al: *APOE* modulates microglial immunometabolism in response to age, amyloid pathology, and inflammatory challenge. Cell reports 2023, 42(3).

111. Rhein V, Song X, Wiesner A, Ittner LM, Baysang G, Meier F, Ozmen L, Bluethmann H, Dröse S, Brandt U et al: Amyloid-β and tau synergistically impair the oxidative phosphorylation system in triple transgenic Alzheimer’s disease mice. Proceedings of the National Academy of Sciences 2009, 106(47):20057–20062.

112. Alldred MJ, Lee SH, Stutzmann GE, Ginsberg SD: Oxidative Phosphorylation Is Dysregulated Within the Basocortical Circuit in a 6-month old Mouse Model of Down Syndrome and Alzheimer’s Disease. Frontiers in Aging Neuroscience 2021, 13.

113. Sharma N, Banerjee R, Davis RL: Early Mitochondrial Defects in the 5xFAD Mouse Model of Alzheimer’s Disease. Journal of Alzheimer’s disease : JAD 2023, 91(4):1323–1338.

114. Naia L, Shimozawa M, Bereczki E, Li X, Liu J, Jiang R, Giraud R, Leal NS, Pinho CM, Berger E et al: Mitochondrial hypermetabolism precedes impaired autophagy and synaptic disorganization in App knock-in Alzheimer mouse models. Mol Psychiatry 2023, 28(9):3966–3981.

115. Lunnon K, Keohane A, Pidsley R, Newhouse S, Riddoch-Contreras J, Thubron EB, Devall M, Soininen H, Kłoszewska I, Mecocci P et al: Mitochondrial genes are altered in blood early in Alzheimer’s disease. Neurobiology of Aging 2017, 53:36–47.

116. Manczak M, Park BS, Jung Y, Reddy PH: Differential expression of oxidative phosphorylation genes in patients with Alzheimer’s disease. NeuroMolecular Medicine 2004, 5(2):147–162.

117. Demetrius LA, Eckert A, Grimm A: Sex differences in Alzheimer&#x2019;s disease: metabolic reprogramming and therapeutic intervention. Trends in Endocrinology & Metabolism 2021, 32(12):963–979.

118. Dave A, Hansen N, Downey R, Johnson C: FDG-PET Imaging of Dementia and Neurodegenerative Disease. *Seminars in Ultrasound*, CT and MRI 2020, 41(6):562–571.

119. Settembre C, Fraldi A, Jahreiss L, Spampanato C, Venturi C, Medina D, de Pablo R, Tacchetti C, Rubinsztein DC, Ballabio A: A block of autophagy in lysosomal storage disorders. Human Molecular Genetics 2008, 17(1):119–129.

120. Villani GRD, Gargiulo N, Faraonio R, Castaldo S, Gonzalez y Reyero E, Di Natale P: Cytokines, neurotrophins, and oxidative stress in brain disease from mucopolysaccharidosis IIIB. Journal of Neuroscience Research 2007, 85(3):612–622.

121. Martins C, Hůlková H, Dridi L, Dormoy-Raclet V, Grigoryeva L, Choi Y, Langford-Smith A, Wilkinson FL, Ohmi K, DiCristo G et al: Neuroinflammation, mitochondrial defects and neurodegeneration in mucopolysaccharidosis III type C mouse model. Brain 2015, 138(Pt 2):336–355.

122. Pará C, Bose P, Bruno L, Freemantle E, Taherzadeh M, Pan X, Han C, McPherson PS, Lacaille J-C, Bonneil É, et al: Early defects in mucopolysaccharidosis type IIIC disrupt excitatory synaptic transmission. JCI Insight 2021, 6(15).

123. Pshezhetsky AV: Crosstalk between 2 organelles: Lysosomal storage of heparan sulfate causes mitochondrial defects and neuronal death in mucopolysaccharidosis III type C. Rare Diseases 2015, 3(1):e1049793.

124. Haust MD: Mitochondrial budding and morphogenesis of cytoplasmic vacuoles in hepatocytes of children with the hurler syndrome and sanfilippo disease. Experimental and Molecular Pathology 1968, 9(2):242–257.

125. Sly WS, Quinton BA, McAlister WH, Rimoin DL: Beta glucuronidase deficiency: Report of clinical, radiologic, and biochemical features of a new mucopolysaccharidosis. The Journal of Pediatrics 1973, 82(2):249–257.

126. Tomatsu S, Gutierrez MA, Ishimaru T, Peña OM, Montaño AM, Maeda H, Velez-Castrillon S, Nishioka T, Fachel AA, Cooper A et al: Heparan sulfate levels in mucopolysaccharidoses and mucolipidoses. Journal of Inherited Metabolic Disease 2005, 28(5):743–757.

127. Parente MK, Rozen R, Seeholzer SH, Wolfe JH: Integrated analysis of proteome and transcriptome changes in the mucopolysaccharidosis type VII mouse hippocampus. Molecular Genetics and Metabolism 2016, 118(1):41–54.

128. Heon-Roberts R, Nguyen ALA, Pshezhetsky AV: Molecular Bases of Neurodegeneration and Cognitive Decline, the Major Burden of Sanfilippo Disease. J Clin Med 2020, 9(2):344.

129. Johnson DE, Ostrowski P, Jaumouillé V, Grinstein S: The position of lysosomes within the cell determines their luminal pH. Journal of Cell Biology 2016, 212(6):677–692.

130. Mindell JA: Lysosomal Acidification Mechanisms. Annual Review of Physiology 2012, 74(1):69–86.

131. Colacurcio DJ, Nixon RA: Disorders of lysosomal acidification-The emerging role of v-ATPase in aging and neurodegenerative disease. Ageing Res Rev 2016, 32:75–88.

132. Mattison KA, Tossing G, Mulroe F, Simmons C, Butler KM, Schreiber A, Alsadah A, Neilson DE, Naess K, Wedell A et al: ATP6V0C variants impair V-ATPase function causing a neurodevelopmental disorder often associated with epilepsy. Brain 2023, 146(4):1357–1372.

133. Kim SH, Cho YS, Kim Y, Park J, Yoo SM, Gwak J, Kim Y, Gwon Y, Kam TI, Jung YK: Endolysosomal impairment by binding of amyloid beta or MAPT/Tau to V-ATPase and rescue via the HYAL-CD44 axis in Alzheimer disease. Autophagy 2023, 19(8):2318–2337.

134. Mangieri LR, Mader BJ, Thomas CE, Taylor CA, Luker AM, Tse TE, Huisingh C, Shacka JJ: ATP6V0C Knockdown in Neuroblastoma Cells Alters Autophagy-Lysosome Pathway Function and Metabolism of Proteins that Accumulate in Neurodegenerative Disease. PLOS ONE 2014, 9(4):e93257.

135. Jiang Y, Sato Y, Im E, Berg M, Bordi M, Darji S, Kumar A, Mohan PS, Bandyopadhyay U, Diaz A et al: Lysosomal Dysfunction in Down Syndrome Is APP-Dependent and Mediated by APP-βCTF (C99). J Neurosci 2019, 39(27):5255–5268.

136. Lee JH, McBrayer MK, Wolfe DM, Haslett LJ, Kumar A, Sato Y, Lie PP, Mohan P, Coffey EE, Kompella U et al: Presenilin 1 Maintains Lysosomal Ca(2+) Homeostasis via TRPML1 by Regulating vATPase-Mediated Lysosome Acidification. Cell reports 2015, 12(9):1430–1444.

137. Lee J-H, Yang D-S, Goulbourne CN, Im E, Stavrides P, Pensalfini A, Chan H, Bouchet-Marquis C, Bleiwas C, Berg MJ et al: Faulty autolysosome acidification in Alzheimer’s disease mouse models induces autophagic build-up of Aβ in neurons, yielding senile plaques. Nature Neuroscience 2022, 25(6):688–701.

138. Prasad H, Rao R: Amyloid clearance defect in ApoE4 astrocytes is reversed by epigenetic correction of endosomal pH. Proc Natl Acad Sci U S A 2018, 115(28):E6640–e6649.

139. Im E, Jiang Y, Stavrides PH, Darji S, Erdjument-Bromage H, Neubert TA, Choi JY, Wegiel J, Lee JH, Nixon RA: Lysosomal dysfunction in Down syndrome and Alzheimer mouse models is caused by v-ATPase inhibition by Tyr(682)-phosphorylated APP βCTF. Sci Adv 2023, 9(30):eadg1925.

140. Fraldi A, Annunziata F, Lombardi A, Kaiser H-J, Medina DL, Spampanato C, Fedele AO, Polishchuk R, Sorrentino NC, Simons K, Ballabio A: Lysosomal fusion and SNARE function are impaired by cholesterol accumulation in lysosomal storage disorders. The EMBO Journal 2010, 29(21):3607–3620.

141. Bach G, Chen C-S, Pagano RE: Elevated lysosomal pH in Mucolipidosis type IV cells. Clinica Chimica Acta 1999, 280(1-2):173–179.

142. Bourdenx M, Daniel J, Genin E, Soria FN, Blanchard-Desce M, Bezard E, Dehay B: Nanoparticles restore lysosomal acidification defects: Implications for Parkinson and other lysosomal-related diseases. Autophagy 2016, 12(3):472–483.

143. Holopainen JM, Saarikoski J, Kinnunen PK, Järvelä I: Elevated lysosomal pH in neuronal ceroid lipofuscinoses (NCLs). European Journal of Biochemistry 2001, 268(22):5851–5856.

144. Pereira VG, Gazarini ML, Rodrigues LC, Da Silva FH, Han SW, Martins AM, Tersariol IL, D’Almeida V: Evidence of lysosomal membrane permeabilization in mucopolysaccharidosis type I: rupture of calcium and proton homeostasis. Journal of cellular physiology 2010, 223(2):335–342.

145. Yambire KF, Rostosky C, Watanabe T, Pacheu-Grau D, Torres-Odio S, Sanchez-Guerrero A, Senderovich O, Meyron-Holtz EG, Milosevic I, Frahm J et al: Impaired lysosomal acidification triggers iron deficiency and inflammation in vivo. eLife 2019, 8:e51031.

146. Sternberg Z, Hu Z, Sternberg D, Waseh S, Quinn JF, Wild K, Jeffrey K, Zhao L, Garrick M: Serum Hepcidin Levels, Iron Dyshomeostasis and Cognitive Loss in Alzheimer’s Disease. Aging and disease 2017, 8(2):215–227.

147. Smith MA, Zhu X, Tabaton M, Liu G, McKeel Jr DW, Cohen ML, Wang X, Siedlak SL, Dwyer BE, Hayashi T et al: Increased Iron and Free Radical Generation in Preclinical Alzheimer Disease and Mild Cognitive Impairment. Journal of Alzheimer’s Disease 2010, 19:363–372.

148. Crespo ÂC, Silva B, Marques L, Marcelino E, Maruta C, Costa S, Timóteo Â, Vilares A, Couto FS, Faustino P et al: Genetic and biochemical markers in patients with Alzheimer’s disease support a concerted systemic iron homeostasis dysregulation. Neurobiology of Aging 2014, 35(4):777–785.

149. Damulina A, Pirpamer L, Soellradl M, Sackl M, Tinauer C, Hofer E, Enzinger C, Gesierich B, Duering M, Ropele S et al: Cross-sectional and Longitudinal Assessment of Brain Iron Level in Alzheimer Disease Using 3-T MRI. Radiology 2020, 296(3):619–626.

150. Kenkhuis B, Somarakis A, de Haan L, Dzyubachyk O, Ijsselsteijn ME, de Miranda NFCC, Lelieveldt BPF, Dijkstra J, van Roon-Mom WMC, Höllt T, van der Weerd L: Iron loading is a prominent feature of activated microglia in Alzheimer’s disease patients. Acta Neuropathologica Communications 2021, 9(1):27.

151. Ayton S, Wang Y, Diouf I, Schneider JA, Brockman J, Morris MC, Bush AI: Brain iron is associated with accelerated cognitive decline in people with Alzheimer pathology. Mol Psychiatry 2020, 25(11):2932–2941.

152. Puy V, Darwiche W, Trudel S, Gomila C, Lony C, Puy L, Lefebvre T, Vitry S, Boullier A, Karim Z, Ausseil J: Predominant role of microglia in brain iron retention in Sanfilippo syndrome, a pediatric neurodegenerative disease. Glia 2018, 66(8):1709–1723.

153. Brady J, Trehan A, Landis D, Toro C: Mucopolysaccharidosis type IIIB (MPS IIIB) masquerading as a behavioural disorder. BMJ case reports 2013, 2013.

154. Ding Q, Markesbery WR, Chen Q, Li F, Keller JN: Ribosome dysfunction is an early event in Alzheimer’s disease. J Neurosci 2005, 25(40):9171–9175.

155. Honda K, Smith MA, Zhu X, Baus D, Merrick WC, Tartakoff AM, Hattier T, Harris PL, Siedlak SL, Fujioka H et al: Ribosomal RNA in Alzheimer Disease Is Oxidized by Bound Redox-active Iron*. Journal of Biological Chemistry 2005, 280(22):20978–20986.

156. Parente MK, Rozen R, Cearley CN, Wolfe JH: Dysregulation of Gene Expression in a Lysosomal Storage Disease Varies between Brain Regions Implicating Unexpected Mechanisms of Neuropathology. PLOS ONE 2012, 7(3):e32419.

157. DiRosario J, Divers E, Wang C, Etter J, Charrier A, Jukkola P, Auer H, Best V, Newsom DL, McCarty DM, Fu H: Innate and adaptive immune activation in the brain of MPS IIIB mouse model. Journal of Neuroscience Research 2009, 87(4):978–990.

158. Taherzadeh M, Zhang E, Londono I, De Leener B, Wang S, Cooper JD, Kennedy TE, Morales CR, Chen Z, Lodygensky GA, Pshezhetsky AV: Severe central nervous system demyelination in Sanfilippo disease. Frontiers in Molecular Neuroscience 2023, 16.

159. Yellajoshyula D, Pappas SS, Rogers AE, Choudhury B, Reed X, Ding J, Cookson MR, Shakkottai VG, Giger RJ, Dauer WT: THAP1 modulates oligodendrocyte maturation by regulating ECM degradation in lysosomes. Proceedings of the National Academy of Sciences 2021, 118(31):e2100862118.

160. Sloane JA, Batt C, Ma Y, Harris ZM, Trapp B, Vartanian T: Hyaluronan blocks oligodendrocyte progenitor maturation and remyelination through TLR2. Proceedings of the National Academy of Sciences 2010, 107(25):11555–11560.

161. Siebert JR, Osterhout DJ: The inhibitory effects of chondroitin sulfate proteoglycans on oligodendrocytes. Journal of Neurochemistry 2011, 119(1):176–188.

162. Tamagawa K, Morimatsu Y, Fujisawa K, Hara A, Taketomi T: Neuropathological study and chemico-pathoiogical correlation in sibling cases of Sanfilippo syndrome type B. Brain and Development 1985, 7(6):599–609.

163. Barone R, Nigro F, Triulzi F, Musumeci S, Fiumara A, Pavone L: Clinical and neuroradiological follow-up in mucopolysaccharidosis type III (Sanfilippo syndrome). Neuropediatrics 1999, 30(5):270–274.

164. Lee S, Viqar F, Zimmerman ME, Narkhede A, Tosto G, Benzinger TLS, Marcus DS, Fagan AM, Goate A, Fox NC et al: White matter hyperintensities are a core feature of Alzheimer’s disease: Evidence from the dominantly inherited Alzheimer network. Annals of Neurology 2016, 79(6):929–939.

165. Prins ND, Scheltens P: White matter hyperintensities, cognitive impairment and dementia: an update. Nature Reviews Neurology 2015, 11(3):157–165.

166. Uhlén M, Fagerberg L, Hallström BM, Lindskog C, Oksvold P, Mardinoglu A, Sivertsson Å, Kampf C, Sjöstedt E, Asplund A et al: Tissue-based map of the human proteome. Science 2015, 347(6220):1260419.

167. Balak CD, Schlachetzki JCM, Lana AJ, West E, Hong C, DuGal J, Zhou Y, Li B, Saisan P, Spann NJ et al: Mechanisms driving epigenetic and transcriptional responses of microglia in a neurodegenerative lysosomal storage disorder model. bioRxiv 2024:2024.2011.2012.623296.

168. Rovira M, Ferrero G, Miserocchi M, Montanari A, Wittamer V: A single-cell transcriptomic atlas reveals resident dendritic-like cells in the zebrafish brain parenchyma. In.: eLife Sciences Publications, Ltd; 2024.

169. Diebold BA, Bokoch GM: Molecular basis for Rac2 regulation of phagocyte NADPH oxidase. Nature Immunology 2001, 2(3):211–215.

170. Zou Y, Xiong J-b, Ma K, Wang A-Z, Qian K-J: Rac2 deficiency attenuates CCl4-induced liver injury through suppressing inflammation and oxidative stress. Biomedicine & Pharmacotherapy 2017, 94:140–149.

171. Lardelli M, Baer L, Hin N, Allen A, Pederson SM, Barthelson K: The Use of Zebrafish in Transcriptome Analysis of the Early Effects of Mutations Causing Early Onset Familial Alzheimer’s Disease and Other Inherited Neurodegenerative Conditions. Journal of Alzheimer’s Disease 2023, Preprint:1-15.

172. Kizil C, Kaslin J, Kroehne V, Brand M: Adult neurogenesis and brain regeneration in zebrafish. Dev Neurobiol 2012, 72(3):429–461.

173. Perez-Riverol Y, Bai J, Bandla C, García-Seisdedos D, Hewapathirana S, Kamatchinathan S, Kundu Deepti J, Prakash A, Frericks-Zipper A, Eisenacher M et al: The PRIDE database resources in 2022: a hub for mass spectrometry-based proteomics evidences. Nucleic Acids Research 2022, 50(D1):D543–D552.

