## Supplemental Figures for "Multi-omics analyses of early-onset familial Alzheimer’s disease and Sanfilippo syndrome zebrafish models reveal commonalities in disease mechanisms"

**Fig.S1: Intra-family transcriptome analysis strategy. A)** A female zebrafish heterozygous for both the *naglu*^A603fs^ (MPS IIIB) and the *psen1*^Q96_K97del^ (EOfAD-like) mutations was mated with a male zebrafish only heterozygous for *naglu*^A603fs^. The subsequent family of ~100 sibling zebrafish larvae was raised in a single petri dish at 28.5 °C. At 7 days post fertilisation (dpf), the entire family was euthanised simultaneously and each larva placed individually in RNAlater solution. After 24 hours, each larva was bisected transversely at the cloaca. The tail end was used for genomic DNA extraction and genotyping, while the remaining head and trunk was used for RNA-seq. **B)** Two different pairs of zebrafish with the same *psen1* and *naglu* mutant genotypes was mated to generate the adult brain analysis families. Entire families were raised until 6 months of age. For RNA-seq (denoted by *), all fish in each family were euthanised, and their entire heads removed and immersed in 6 volumes of RNAlater solution, and stored at 4°C overnight. Brains were subsequently dissected out of the RNAlater-preserved heads while their corresponding tails were used for genotyping by PCR. For proteomics, the brains were dissected from the skull, snap frozen and stored at -80°C until use.

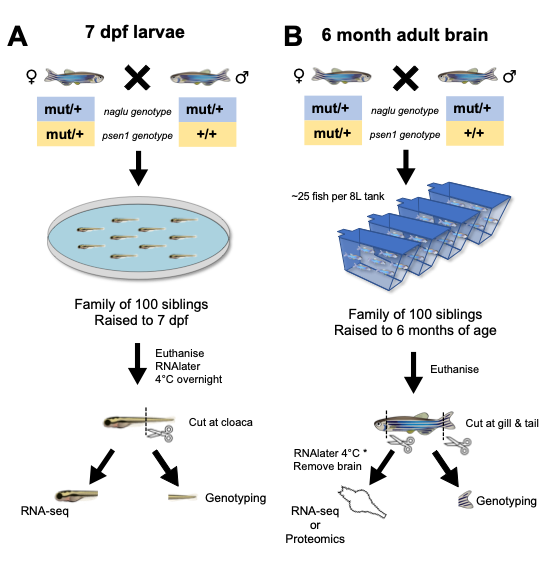

**Fig.S2:** **PCR genotyping of *psen1^Q96_K97del^* in zebrafish***.* **A)** Schematic of a PCR amplifying across the *psen1^Q96_K97^* region in exon 3 of zebrafish *psen1.* The 6 bp deletion of the *psen1^Q96_K97del^* mutation (red) results in a smaller PCR product. **B)** Example 3% agarose in TAE agarose gel electrophoresis showing the wild type and *psen1^Q96_K97del^*/+ mutant PCR products resolved after electrophoresis for 2 hours at 90 volts. The M lane contains the HAPE DNA ladder [1], while positive control lanes contain PCR products from known heterozygous (+/-) and wild type (+/+) zebrafish genomic DNA samples.

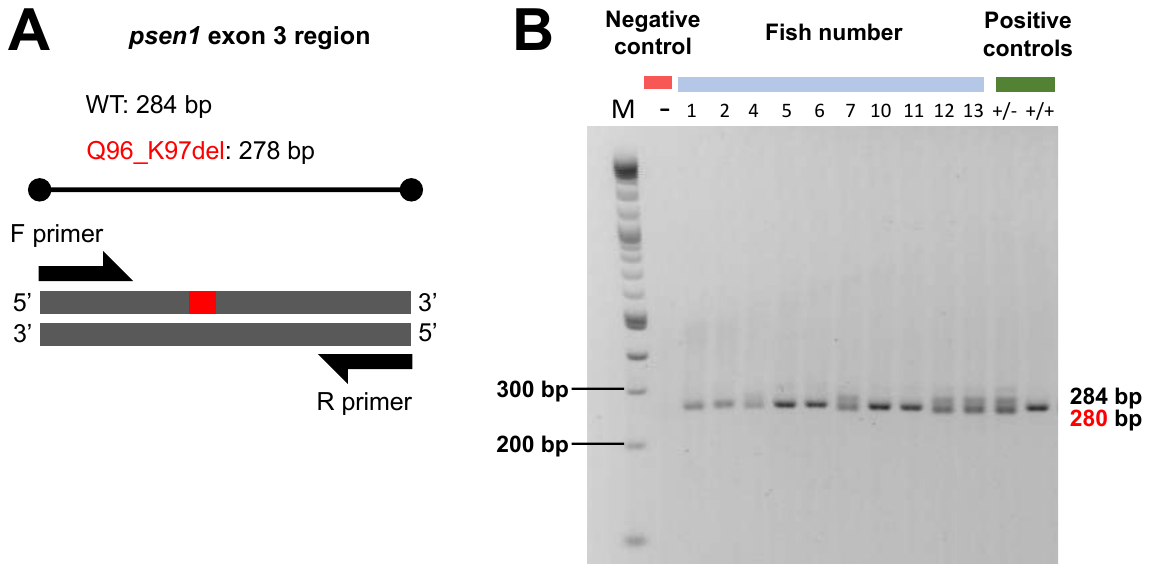

**Fig.S3:** **PCR genotyping of *naglu^A603fs^* in zebrafish***.* **A)** Schematic of a PCR amplifying across the region of the *naglu^A603^* mutation in exon 7 of zebrafish *naglu.* The 6 bp deletion of the *psen1^Q96_K97del^* mutation (red) results in a smaller PCR product. **B)** Example 3% agarose in TAE agarose gel electrophoresis showing the wild type and *psen1^Q96_K97del^*/+ mutant PCR products resolved after 2 hours of electrophoresis at 90 volts. The M lane contains the HAPE DNA ladder [1], and positive control lanes contain PCR products from known heterozygous (+/-) and wild type (+/+) zebrafish genomic DNA samples.

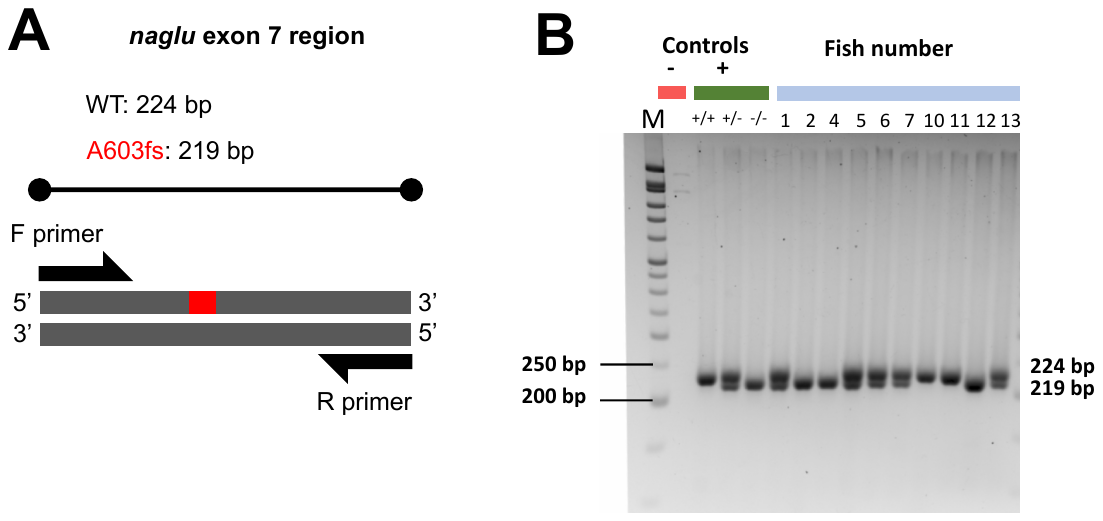

**Fig.S4: Workflow of the snakemake pipeline used for RNA-seq analysis**. Raw reads from each of the lanes were first merged using the *cat* function. The merged, raw reads were trimmed and filtered for quality and length using *fastp* [2]. The resulting reads were aligned to the zebrafish genome (GRCz11, Ensembl release 101) using *STAR* [3]. PCR duplicates were deduplicated using *umi-tools* [4]. Read counts were generated using *featureCounts* [5], and quality of the raw and processed data were assessed using *FastQC* [6] and *ngsReports* [7].

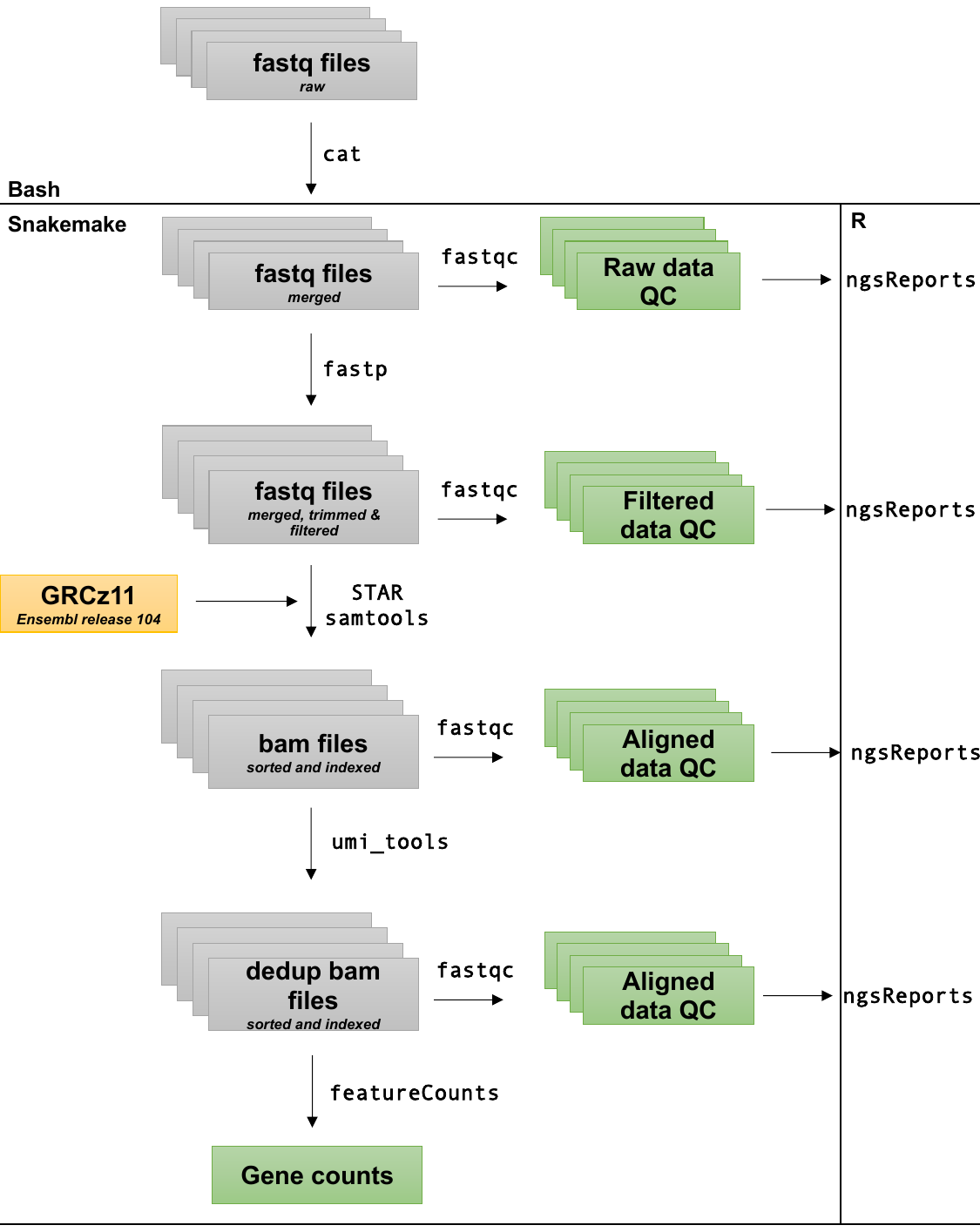

**Fig.S5**: Observed bias for differential expression of genes with their %GC and length in **A)** 7 dpf zebrafish larvae and **B)** 6-month-old adult zebrafish brains. A ranking statistic per gene was calculated as the sign of the logFC (1 for upregulated and -1 for downregulated) multiplied by the negative log_10_ of the p-value from the likelihood ratio tests in edgeR in each mutant. This was plotted against a weighted (by transcript length) average %GC content per gene and average transcript length. The blue generalised additive model fit (gam) lines are not centred on 0, particularly in 6 month (6m) adult brains, indicating a bias. The y-axes in these plots are constrained to be between -5 and 5 for visualisation purposes.

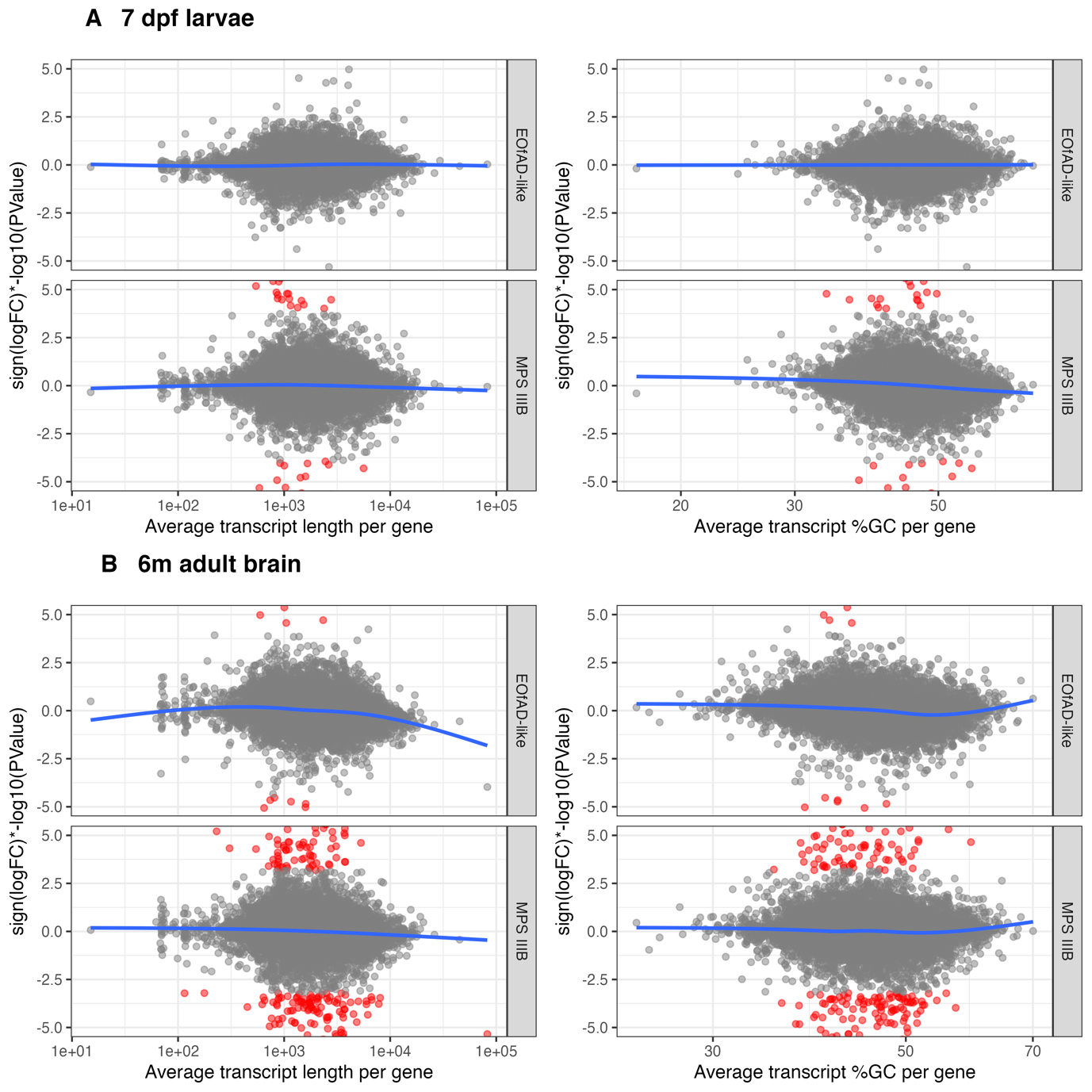

**Fig.S6**: Heatmaps showing the logFC values of genes in the KEGG [8] gene sets for **A)** *lysosome*, **B)** *glycosaminoglycan degradation* and **C)** *other glycan degradation* in 7 dpf EOfAD-like and MPS IIIB zebrafish larvae.

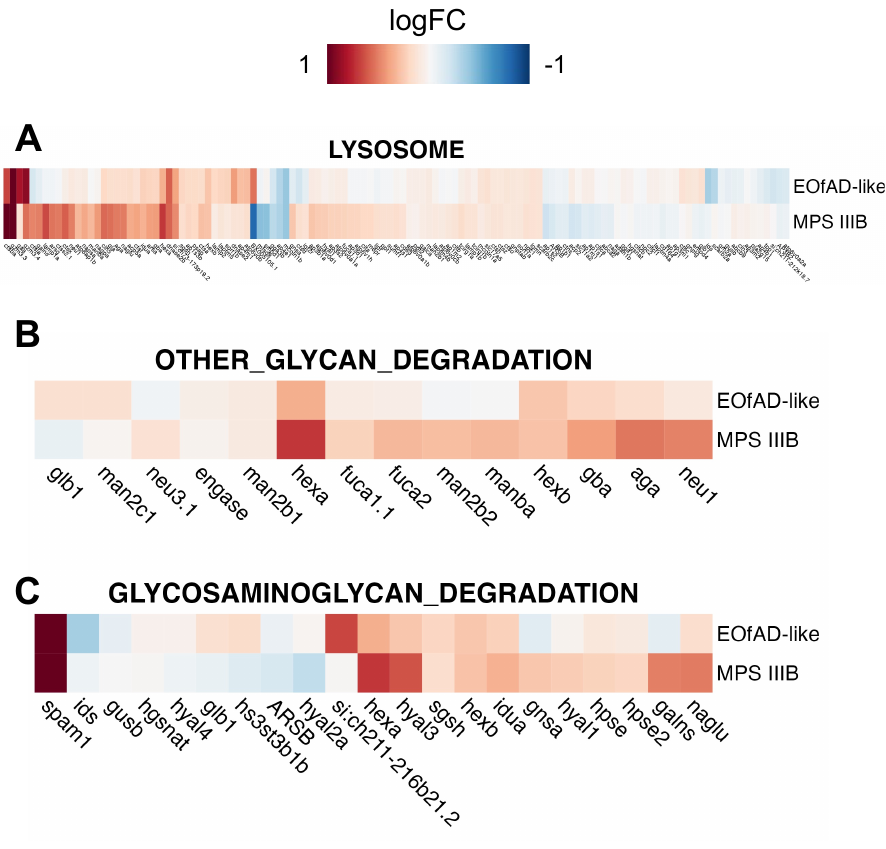

**Fig.S7:** **Changes to ECM gene expression in zebrafish larvae. A)** Pathview [9] visualisation of the KEGG gene set for ECM receptor interaction in EOfAD-like and MPS IIIB zebrafish larvae at 7 dpf. Genes (or groups of genes with similar function) are represented as rectangles, metabolites as circles, and interactions as lines with arrowheads. The genes are coloured by their log_2_FC value in EOfAD-like (left half of rectangle) and MPS IIIB (right half of rectangle) zebrafish larvae. Upregulated genes appear red, downregulated genes appear blue, unchanged elements in grey, and genes considered undetectable in the RNA-seq experiment appear white. This KEGG diagram is modified with permission from KEGG. **B)** Heatmap showing the logFC values for collagen encoding genes in MPS IIIB and zebrafish larvae.

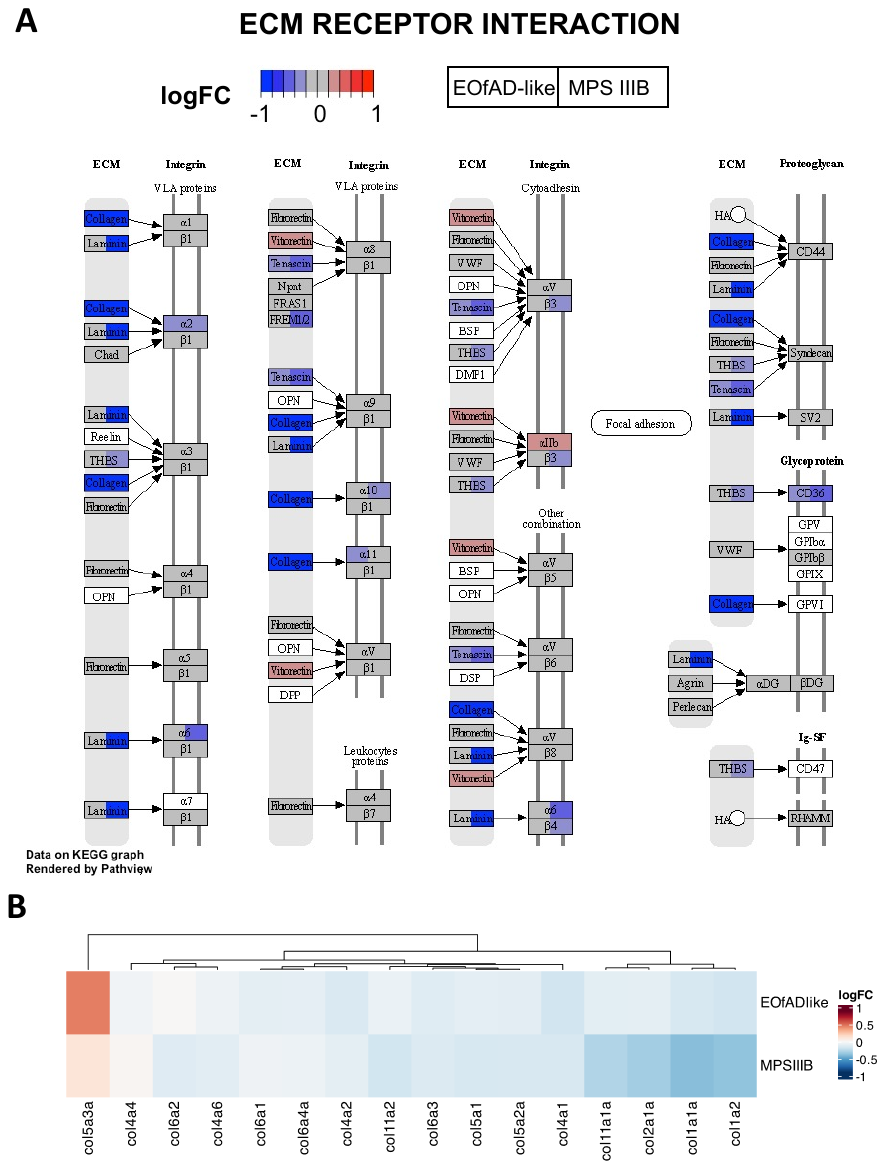

**Fig.S8: Post-hoc power calculation in zebrafish RNA-seq data.** The black line indicates the level of power achieved for a given sample size (n) in **A)** zebrafish larvae and **B)** 6 month old brains. The red dashed line in A) indicates that, when power = 70% (an acceptable value for RNA-seq), n = 20 individual larvae are required. The blue dashed line indicates approximately 50% power was achieved in the 7 dpf dataset at n = 8. Approximately 80% power was achieved at n = 8 in adult brains (blue dashed line).

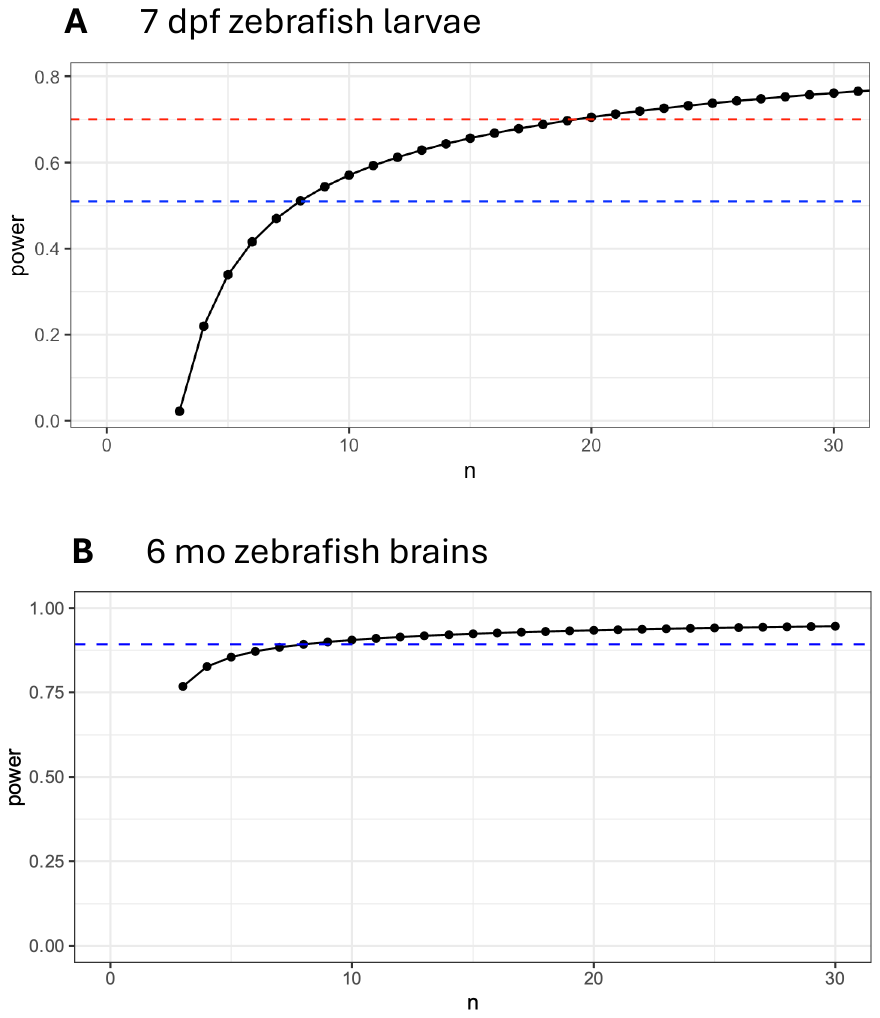

**Fig.S9: Heatmap showing of shared leading-edge genes in MPS IIIB and EOfAD-like brain transcriptomes.** The logFC values for shared leading-edge genes from the **A)** ribosome, **B)** oxidative phosphorylation, and **C)** lysosome gene sets are shown. Rows represent pairwise contrasts between mutant genotypes (by sex) to their wild-type siblings, while columns represent individual genes. The colour scale indicates logFC values, with red representing upregulated genes and blue indicating downregulated genes. M: male, F: female.

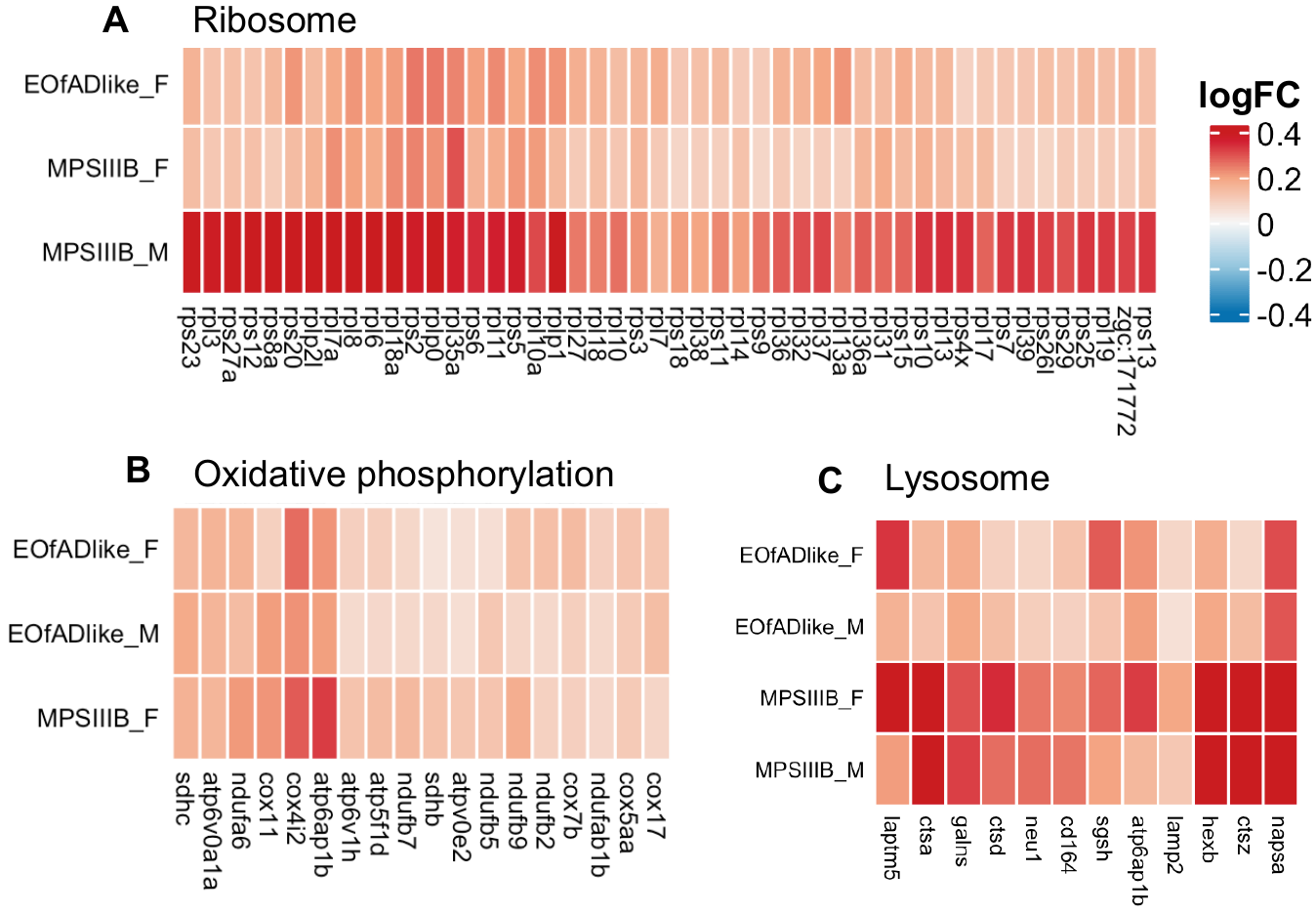

**Fig.S10: STRING analysis of DE proteins in MPS IIIB brain proteomes.** Each node represents a DE protein in **A)** male or **B)** female MPS IIIB brain proteomes relative to their wild type siblings. Nodes were clustered using k-means, setting k = 7 and clusters containing more than 2 proteins are annotated based on their over-represented GO terms. Only connected nodes are displayed for visualisation purposes. Nodes are connected by weighted edges by the confidence of the interaction (thicker nodes mean higher confidence). Protein-protein interactions were set to "high confidence" (0.7) based on STRING criteria.

**Fig.S11: Heatmap showing the shared leading edge proteins in MPS III and EOfAD-like zebrafish brain proteomes.** The logFC values for shared leading-edge proteins from the **A)** lysosome and **B)** ribosome protein sets are shown. Rows represent pairwise contrasts between mutant genotypes (by sex) to their wild-type siblings, while columns represent individual genes. The colour scale indicates logFC values, with red representing upregulated genes and blue indicating downregulated proteins. M: male, F: female.

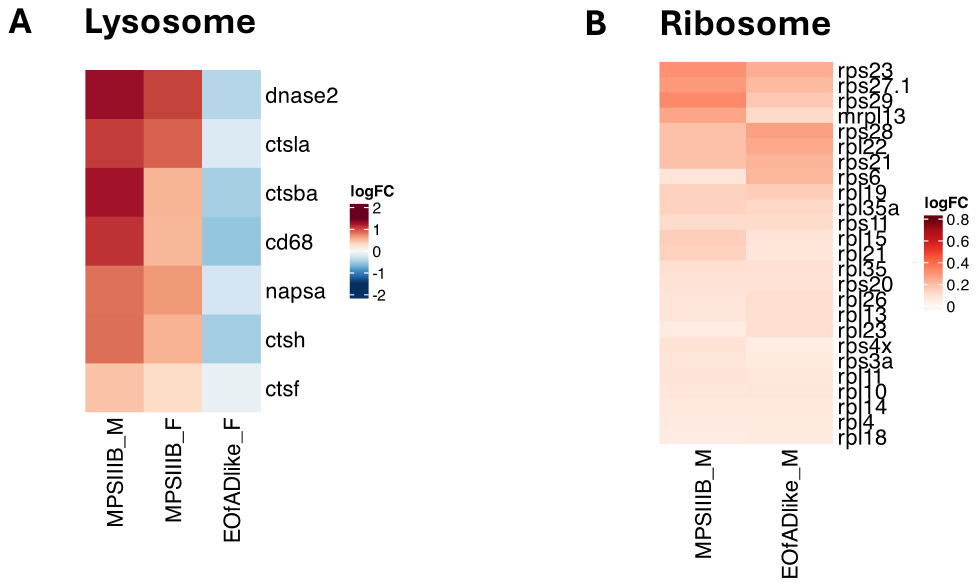

**Fig.S12: Heatmap of the logFC of genes and proteins in the oxidative phosphorylation pathway in MPS IIIB and EOfAD-like zebrafish brains.** Rows represent pairwise contrasts between mutant genotypes (by sex) to their wild-type siblings, while columns represent individual proteins. The columns are grouped by the complex they belong to in the electron transport chain. The colour scale indicates logFC values, with red representing upregulated genes and blue indicating downregulated proteins. M: male, F: female

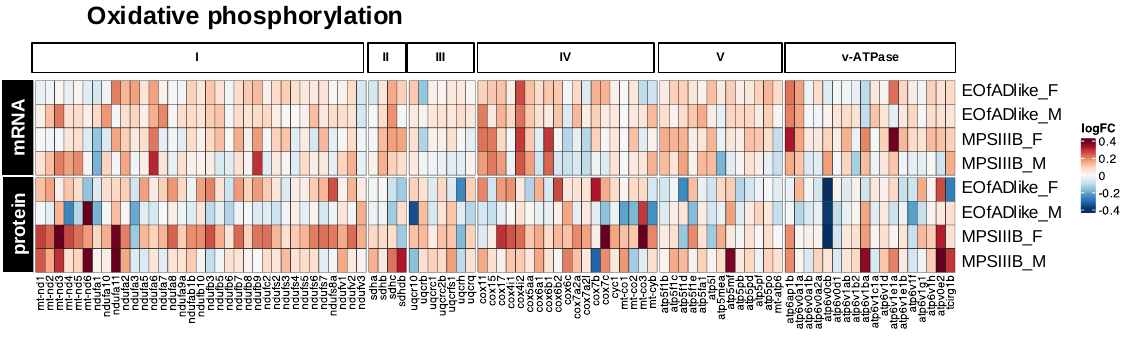

**Fig.S13: Heatmap of the logFC of genes and proteins in the immune-related pathway in MPS IIIB and EOfAD-like zebrafish brains.** Rows represent pairwise contrasts between mutant genotypes (by sex) to their wild-type siblings, while columns represent individual proteins. The colour scale indicates logFC values, with red representing upregulated genes and blue indicating downregulated proteins in **A)** *Fc gamma r mediated phagocytosis* and **B)** *natural killer cell mediated cytotoxicity*. Grey indicates the gene or protein was not detected in the respective experiment. M: male, F: female

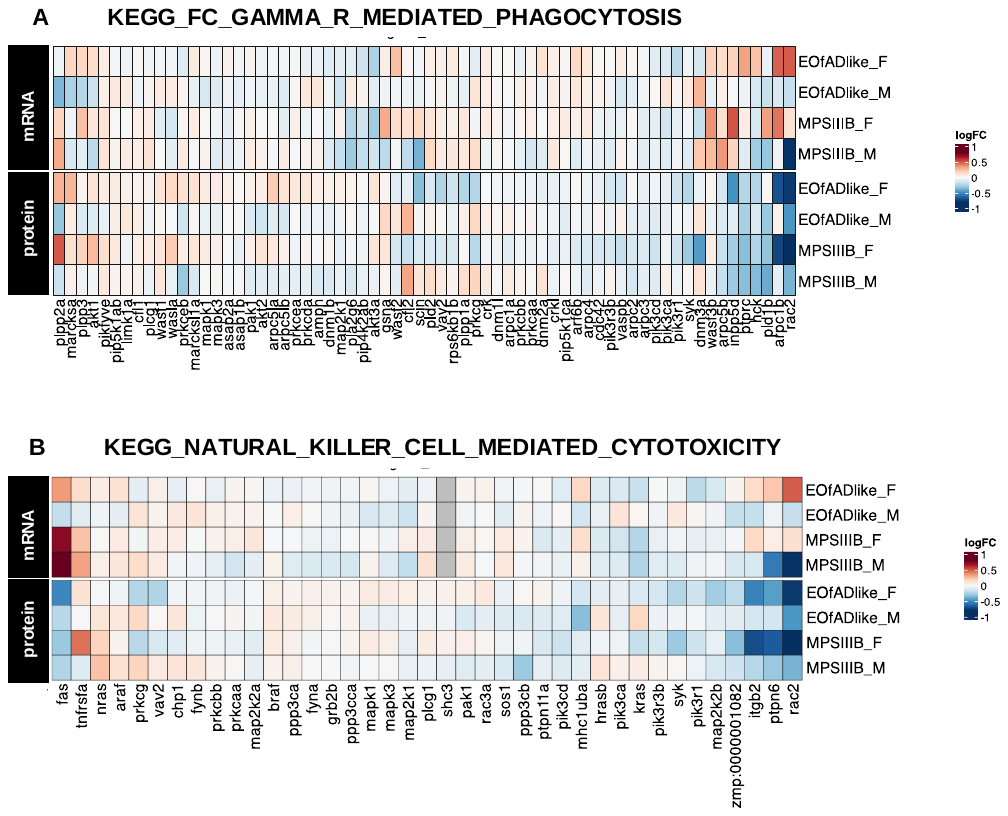

**Fig.S14**: Heatmap showing the enrichment analysis results for iron-responsive element (ire)-containing genes in 7 days post-fertilisation (dpf) zebrafish larvae, and 6-month-old (6 mo) zebrafish brain samples. Each cell in the heatmap represents an IRE (ire) gene set which is coloured based on the -log_10_ of the harmonic mean p-value, with brighter colours indicating more significant results. The numbers inside the cells in the heatmap are the FDR-adjusted harmonic mean p-values. The gene sets are labelled with “hq” if the IRE gene set only contains those genes with consensus IREs. The gene sets are labelled “all” if they contain any consensus and non-consensus ire-containing genes. The “5” and “3” in the IRE gene set labels indicate if the genes encode transcripts with an IRE(s) in the 5’ or 3’ untranslated region.

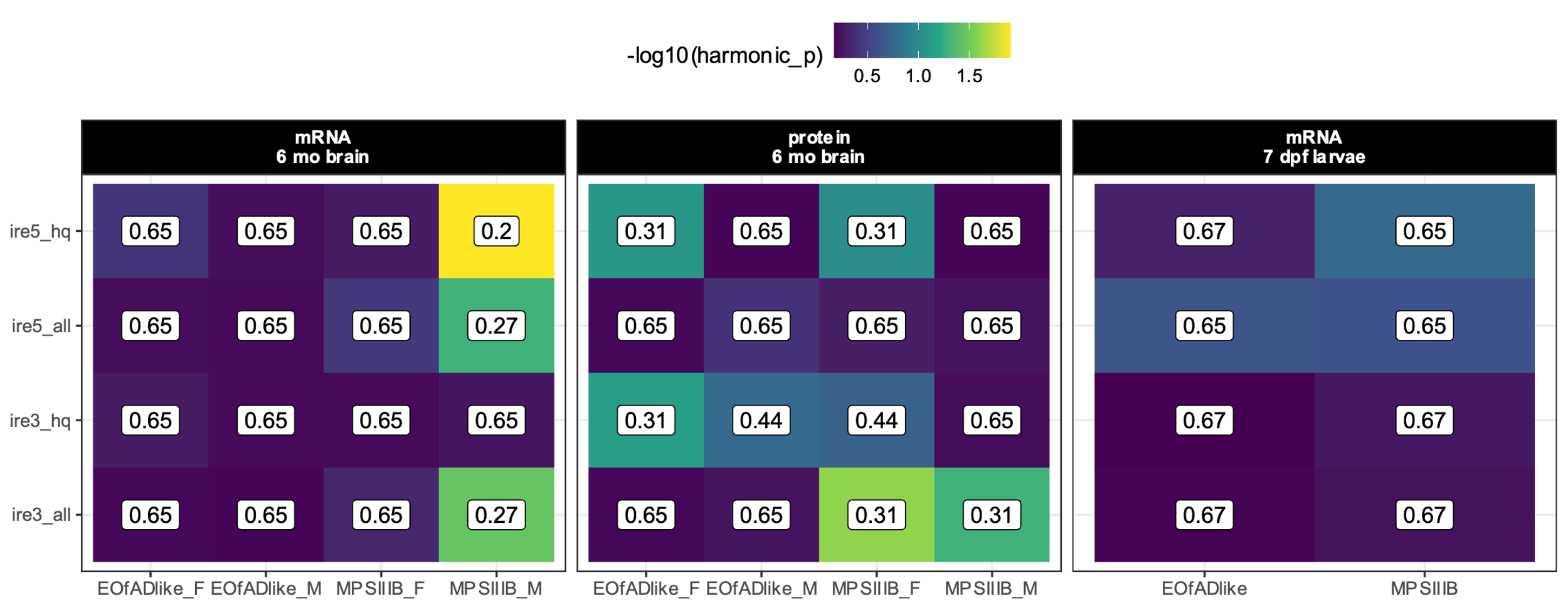

**Fig.S15: KEGG gene sets significantly altered in pools of EOfAD-like mutant zebrafish at 7 dpf.** Only the KEGG gene sets reaching the FDR-adjusted, harmonic mean p-value of < 0.05 are shown. The plot shows the -log_10_ of the harmonic mean p values for each KEGG gene set, with the gene sets ordered according to their harmonic mean p-value. The fill colour indicates the significance level of each gene set, with brighter, more yellow colours representing higher significance.

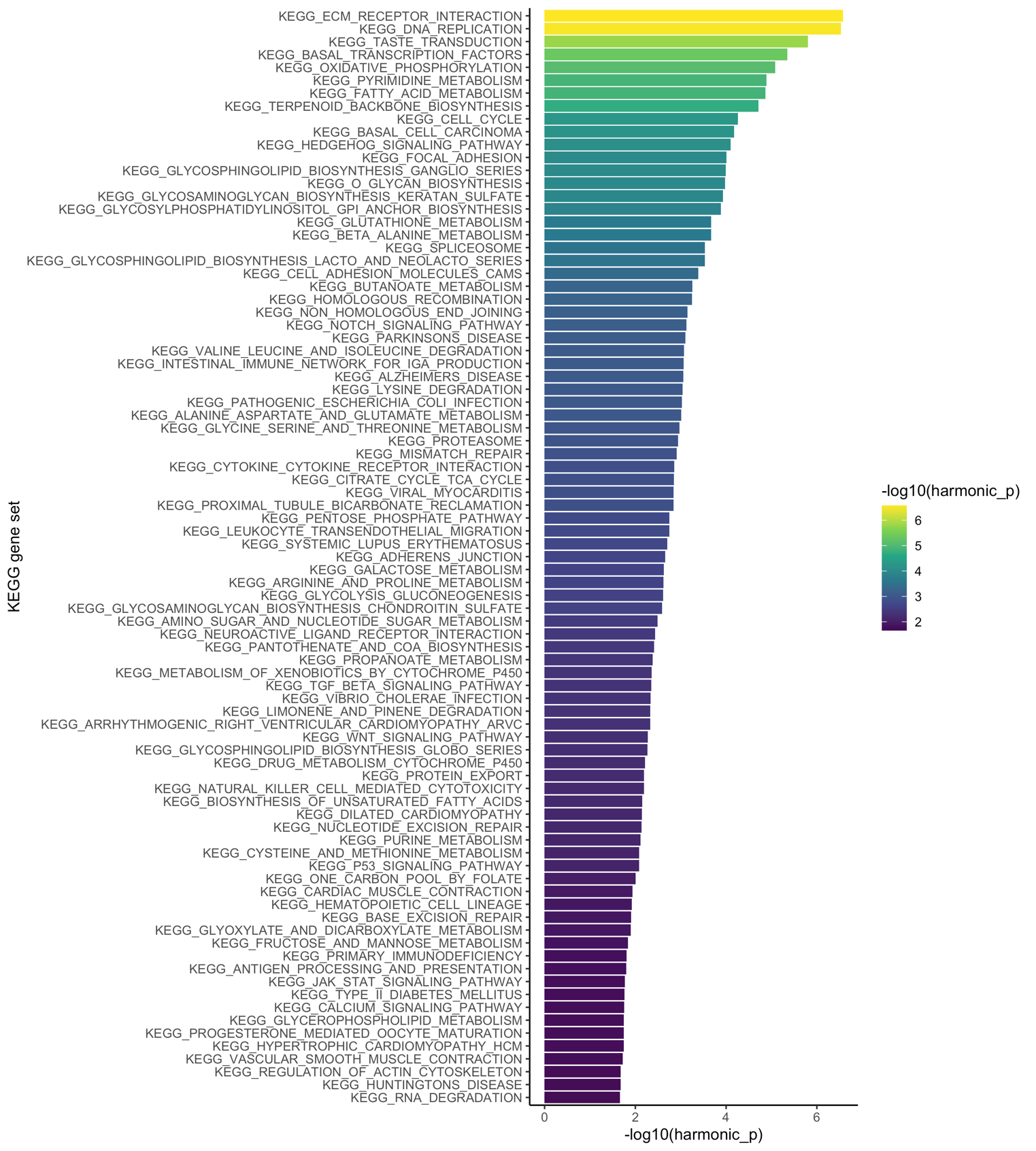

References for supplemental figures:

1.	Allen, A.G., K. Barthelson, and M. Lardelli, *pHAPE: a plasmid for production of DNA size marker ladders for gel electrophoresis.* Biology Methods and Protocols, 2023. **8**(1): p. bpad015.

2.	Chen, S., et al., *fastp: an ultra-fast all-in-one FASTQ preprocessor.* Bioinformatics, 2018. **34**(17): p. i884-i890.

3.	Dobin, A., et al., *STAR: ultrafast universal RNA-seq aligner.* Bioinformatics, 2013. **29**(1): p. 15-21.

4.	Smith, T.S., A. Heger, and I. Sudbery, *UMI-tools: Modelling sequencing errors in Unique Molecular Identifiers to improve quantification accuracy.* Genome Research, 2017.

5.	Liao, Y., G.K. Smyth, and W. Shi, *featureCounts: an efficient general purpose program for assigning sequence reads to genomic features.* Bioinformatics, 2014. **30**(7): p. 923-930.

6.	Andrews, S., *FastQC: a quality control tool for high throughput sequence data*. 2010, Babraham Bioinformatics, Babraham Institute, Cambridge, United Kingdom.

7.	Ward, C.M., T.H. To, and S.M. Pederson, *ngsReports: a Bioconductor package for managing FastQC reports and other NGS related log files.* Bioinformatics, 2020. **36**(8): p. 2587-2588.

8.	Kanehisa, M. and S. Goto, *KEGG: Kyoto Encyclopedia of Genes and Genomes.* Nucleic Acids Research, 2000. **28**(1): p. 27-30.

9.	Luo, W. and C. Brouwer, *Pathview: an R/Bioconductor package for pathway-based data integration and visualization.* Bioinformatics, 2013. **29**(14): p. 1830-1831.
